# The DiffInvex evolutionary model for conditional somatic selection identifies chemotherapy resistance genes in 10,000 cancer genomes

**DOI:** 10.1101/2024.06.17.599362

**Authors:** Ahmed Khalil, Fran Supek

## Abstract

Tumors often show an initial response to chemotherapy, but then develop resistance, leading to relapse and poor prognosis. We hypothesized that a genomic comparison of mutations in pre-treated versus treatment-naive tumors would serve to identify genes that confer resistance. A challenge in such an analysis is that therapy alters mutation burdens and signatures, confounding association studies and complicating identifying causal, selected mutations. We developed DiffInvex, a framework for identifying changes in selection acting on individual genes in somatic genomes. Crucially, DiffInvex draws on a mutation rate baseline that accounts for these shifts in neutral mutagenesis during cancer evolution. We applied DiffInvex to 9,953 cancer whole-genomes from 29 cancer types from 8 studies, containing both WGS of treatment-naive tumors and tumors pre-treated by various drugs, identifying genes where point mutations are under conditional positive or negative selection for a certain chemotherapeutic, suggesting resistance mechanisms occurring via point mutation. DiffInvex confirmed well-known chemoresistance-driver mutations in *EGFR*, *ESR1*, *KIT* and *AR* genes as being under conditional positive selection, with additional cancer types identified for *EGFR* and *KIT*. Additionally, DiffInvex identified 11 genes with treatment-associated selection for different classes of therapeutics. In most cases, these genes were common cancer genes including *PIK3CA*, *APC*, *MAP2K4* and *MAP3K1*. This suggests that tumor resistance to therapy via mutation often occurs via selective advantages conferred by known driver genes, rather than via mutations in specialized resistance genes. Various gene-chemotherapy associations were further supported in tests for functional impact of mutations, again implemented in a conditional selection setting, as well as replicating in independent panel or exome sequencing data. In addition to nominating drug resistance genes that could be targeted by future therapeutics, DiffInvex can also be applied to diverse analysis in cancer evolution, such as comparing normal and tumoral tissues, or analyzing subclonal evolution, identifying changes in selection over time.

## Introduction

Chemotherapy is a standard treatment in cancer. It may be used alone or in combination with other treatments such as immunotherapy or radiotherapy to kill rapidly dividing cancer cells. However, the development of resistance to chemotherapy is a rule rather than an exception, and poses a significant challenge in cancer treatment. Many tumors show good initial response to chemotherapy, but then stop responding and relapse^1,2^, suggesting that some genetic or epigenetic features have evolved in the tumor cells to grant them chemotherapy resistance. Identifying these changes and understanding the underlying mechanisms is the first step to countering the resistance to chemotherapy.

Conducting a genomic analysis to compare mutations present in pre-treated tumors with those in treatment-naive tumor genomes that have not yet been treated would enable the identification of genes associated with the emergence of resistance, thereby providing insights into the mechanisms driving treatment resistance in cancer. Over the last decade many large-scale tumor profiling studies using next generation sequencing (NGS), such as Hartwig^3^, PCAWG^4^, POG^5^, DECIDER^6^, GENIE^7^, and FH-FMI CGDB^8^, have provided the genomic mutational landscape of thousands patients before and/or after treatment with heterogeneous combination of drugs. These genomic profiles along with the clinical information enable us to study the tumor evolution during treatment and identify genetic elements that underlie patients’ response to different chemotherapy drugs.

A major challenge in such comparative analysis is that mutation burdens and signatures are altered due to chemotherapy exposure itself, as well as other factors that may differ between the treated and untreated cohorts, both biological and technical. This confounds association studies and complicates identifying causal selected mutations that drive chemotherapy resistance. In other words, observing an enrichment of mutations in a certain gene in pre-treated versus untreated tumors does not necessarily mean these mutations were selected, as the enrichment may have resulted from the treatment’s mutagenic effects or from other confounding. Given the increasing availability of WGS data, mutations in non-coding regions serve as a valuable proxy for tracking neutral changes in coding mutations and estimating the mutation spectrum and the local neutral mutation rate. This provides an empirical baseline that obviates the need to rely on inferring mutation risk from covariate information, such as replication time or gene expression as proxies for mutation rates.

Motivated by the above, we have developed the DiffInvex approach to quantify conditional selection in cancer by comparing selective pressures upon mutations in genes between two or more conditions, while being able to control for arbitrary confounding factors, and drawing on empirical mutation rate baselines in WGS data. While DiffInvex is broadly applicable to diverse comparisons in cancer evolutionary studies, here we apply it to identify driver genes that increase or decrease selection upon chemotherapy treatment using WGS data. We show that DiffInvex could accurately estimate the local mutation rate baseline using the intronic and intergenic regions mutations, comparing favorably to covariate-based baseline. Trinucleotide and pentanucleotide composition and DNA sequence mappability are stringently accounted for via a locus sampling approach. We applied DiffInvex to a large set of nearly 10,000 cancer whole-genomes of 29 cancer types from 9 studies, containing both WGS from treatment-naïve tumors and tumors pre-treated by various chemotherapy drugs, usually administered in combination. DiffInvex confirmed known examples of drug-resistance driver genes, which were seen to be relevant to additional tissues than anticipated. Moreover, DiffInvex identified 11 putative resistance genes for different classes of therapeutics, which were further validated using a test for differential functional impact of the mutations before and after treatment. Importantly, the validated hits in our analysis were largely not novel genes associated with chemoresistance via gaining mutations. Instead, we identify various examples of common cancer genes such as *PIK3CA* or *APC*, in which driver mutations predicted resistance to specific chemotherapies.

## Results

### 8,591 whole-genome sequenced samples from pre-treated and treatment-naive patients

To identify the genes under differential positive selection associated with chemotherapy treatment, we collected the somatic SNV mutations from WGS of 9,953 tumor samples from the cohorts with pre-treated samples of Hartwig Medical Foundation (henceforth “Hartwig”), POG570, DECIDER, and MMRF-COMMPASS (henceforth: MMRF), and the largely treatment-naive WGS from the PCAWG, CPTAC-3, Mutographs, ICGC (other than PCAWG), and OVCARE cohorts. After excluding samples with low-quality data (e.g. samples with tumor purity < 20%, PCAWG- blacklisted samples and MSI samples) or with missing treatment information, and tumor types with less than 5 treated or less than 5 treatment-naive patients, a total of 8,591 tumor samples (3,360 Hartwig, 1,865 PCAWG, and 880 MMRF, 778 ICGC, 541 Mutographs, 510 POG570, 322 CPTAC-3, 202 DECIDER, 133 OVCARE) from 29 tumor types were used in the analyses (see Methods, Supplementary Table 1).This includes 4138 (48%) primary tumor samples and 4453 (52%) metastatic tumor samples (Supplementary Fig. 1a). Among them, 2,730 (32%) samples were obtained from biopsy pre-treated with chemotherapy drug(s) and/or other drug types, and 5,861 (68%) samples were treatment-naive (Supplementary Fig. 1b). Pre-treated samples are from Hartwig (2167 samples), POG570 (424 samples), DECIDER (70 samples) and MMRF (69 samples). Eight out of 29 tumor types have more than 500 samples each (Fig. 1a): breast cancer (BRCA, 1365), prostate cancer (PRAD, 932), multiple myeloma (MM, 882 samples), colorectal cancer (COREAD, 681), esophageal cancer (ESCA, 650), ovarian cancer (OV, 596), non-small cell lung cancer (NSCLC, 559) and pancreatic cancer (PAAD, 525). The Hartwig, PCAWG, and POG570 studies included samples from diverse tumor types (>= 19), while other studies focused on few tumor types (Supplementary Fig. 1c), four tumor types from CPTAC-3, two tumor types from MUTOGRAPHS and ICGC each, and one tumor type from each of DECIDER, OVCARE, and MMRF.

**Figure 1.**
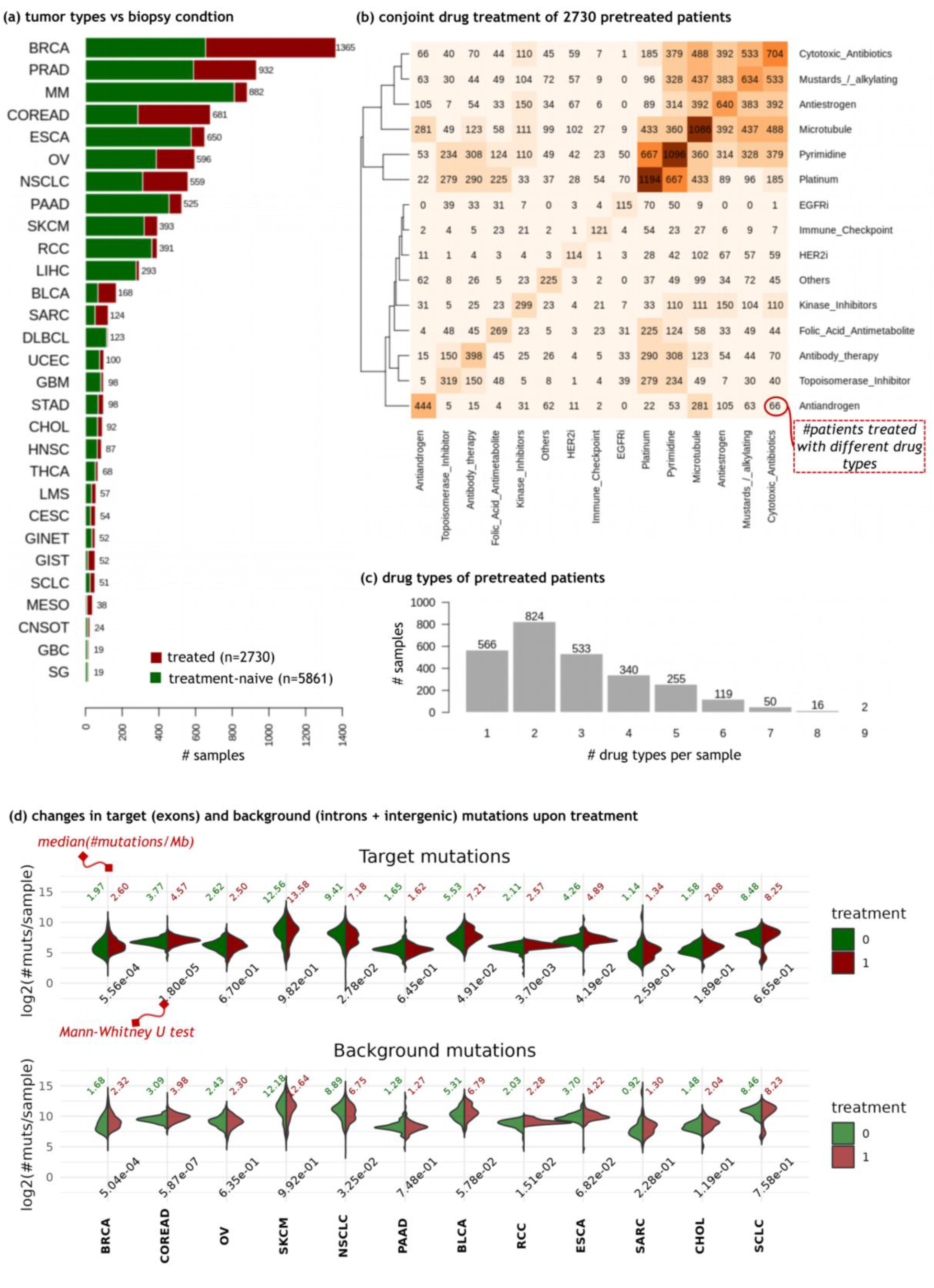
Overview of dataset of pre-treated tumor genomes showing drug combinations and differences in tumor mutation burden. a) Number of treatment-naive and pre-treated patients from the 29 cancer types included in this study. **b)** Number of patients pre-treated with different drug combinations. The elements on the diagonal represent the total number of patients pre-treated with the drug type on the x-axis, with or without other drug types. **c)** Number of drug types that were administered to patients prior to biopsy. **d)** Differences in the tumor mutation burden (here, number of point mutations per sample) between the treatment-naive (in green) and pre-treated (in red) metastatic tumors in different cancer types. The top and bottom panels represent the changes in the exonic (DiffInvex target) regions and intronic (DiffInvex background) regions per sample, respectively. The median mutation burdens per megabase were shown on top of each panel for treatment-naive tumors (in green) and the pre-treated (in red). The numbers on bottom of each panel are the p-values of the Mann-Whitney U test, two-tailed, comparing the tumor mutation burden between the treatment-naive and pre-treated patients.

### Statistics of drug treatments across cancer types show common overlaps in drug regimens

We then examined and classified the drugs that were given to the 2730 patients prior to the biopsy, based on their drugs mechanism-of-action category (https://go.drugbank.com/) dividing them into 15 drug type groups (Fig. 1b, Supplementary Table 2). These included 6 DNA damaging chemotherapy groups (platinum-based alkylating agents, pyrimidine analogs, cytotoxic antibiotics, nitrogen mustards and other alkylating, topoisomerase inhibitor, and folic acid antimetabolites) and 9 other chemotherapy drug groups (microtubule agents, antiestrogens, antiandrogens, antibody therapy, kinase inhibitors, immune checkpoint inhibitors, EGFRi, HER2i, and other drugs).

We observed a wide diversity in the drugs administered to patients in the analyzed data, covering 138 different drugs in total (Supplementary Fig. 1d-e), with most common being platinum-based alkylating, pyrimidine analogs, and microtubule drugs (> 1000 patients each). Most drug types were used across many tumor types: 12 drug types were given to >=5 patients from more than 5 tumor types, while only 3 drug types (antiandrogens, HER2i and EGFRi) were mainly given to one or two tumor types (Supplementary Fig. 1d). This suggests that performing a pan-cancer study would help to boost the power for statistical tests of association between drug exposure and driver mutation.

Most patients were treated with drug combinations of different drug types, while only 548/2730 patients were treated with a single drug type (Fig. 1c). For example, antiestrogen drugs are jointly used with diverse chemotherapy drug groups such as pyrimidine analogs, microtubule agents and cytotoxic antibiotic drugs, while antiandrogen drugs were mainly used with microtubule drugs. These conjoint therapy regimes may confound association studies between individual drugs and the occurrence of mutations in their putative resistance genes, and should thus be rigorously controlled for.

### Controlling for tumor mutation burden changes associated with treatment in tests for selection

Tumor cell mutation burden increases upon treatment with some chemotherapies, and the trinucleotide mutational spectrum typically changes^9^; mutagenicity is clear for the commonly-applied platinum and 5-FU therapies^10,11^. To confirm these trends in our data, we compared the tumor mutation burden (TMB) of point mutations between metastatic tumors in pre-treated versus treatment-naive patients. Expectedly, we found that exonic mutation burden per sample was positively associated with treatment for many tumor types (p-values of 1.8e^-5^, 5.6e^-4^, 3.7e^-3^, 2.8e^- 2^, 4.2e^-2^ and 4.9e^-2^ for COREAD, BRCA, RCC, NSCLC, ESCA and BLCA tumors, respectively, using Mann-Whitney U test) (Fig. 1d). We also observed similar differences in metastatic tumors treated with one drug type or other drug types (Supplementary Fig. 2). For example, comparing tumors treated with platinum drugs versus other drugs showed that the TMBs were significantly different for BRCA, COREAD and NSCLC tumors with p = 2.99e^-02^, 6.41e^-03^, and 6.74e^-06^, respectively. Similarly, the TMBs were significantly altered between patients treated with pyrimidine analogs (this group includes 5-FU and its prodrug capecitabine) and patients treated with other drug types (p = 3.42e^-02^ and 3.34e^-02^ for NSCLC and OV tumors respectively using Mann-Whitney U test). These associations of treatment exposure with exonic mutation burdens - largely consisting of passenger mutations - have potential to significantly confound identification of drug resistance driver mutations.

An opportunity presents itself in rapidly increasing availability of WGS data, which allows to utilize the mutations in the intronic and, optionally, neighboring intergenic regions as a background to model the shifts of exonic mutation burdens. This is the rationale underlying the previous InVEx (introns-versus-exons) method for identifying positive somatic selection in WGS^12^, and here we propose to use the intronic mutations as baseline in a test for differential somatic selection i.e. identifying conditional driver genes. To validate this concept of intronic rate baselines, we repeated the TMB comparisons using the background intronic/neighboring intergenic mutations (see Methods). The treatment-associated changes in the background mutation rates per sample indeed mirrored the pattern of coding exonic mutation changes per sample for most tumor types (Fig. 1d and Supplementary Fig. 3). In particular, the background mutation burden was increased significantly for COREAD, BRCA, RCC and NSCLC metastatic tumors upon treatment (all p values < 0.05, Mann-Whitney U test) (fig. 1d). Similarly, the increase in background mutation burden was significant for platinum drugs versus other drugs for BRCA, COREAD, NSCLC and SARC tumors, and for pyrimidine drugs versus other drugs for NSCLC and OV tumors (p-values < 0.05, Mann-Whitney U test) (Supplementary Fig. 2).

Next, we further examined the differences between drug treatments on the target and background TMB. We observed strong variation of the target (exonic) mutation rate per sample between different drug combinations, normalized by the median mutation rate (Supplementary Fig. 3a). However, these variations diminished when target mutation rate per sample was normalized by the background mutation rate per sample (Supplementary Fig. 3b). Taken together, it is essential to account for deviation in passenger mutation rate upon treatment with different drugs while identifying drug-resistance driver gene associations (i.e. conditional positive selection associated with drug treatment), and intronic and neighboring intergenic mutations could compensate for these deviations.

### DiffInvex framework for quantifying conditional selection in cancer

Motivated by our search for genes under stronger positive selection after chemotherapy exposure, but We have developed DiffInvex (Differential Introns Versus Exons), a statistical method to identify conditional selection on point mutation in cancer (Fig. 2a). Inspired by the previous “InVEx’’ approach^12^, DiffInvex first estimates the local baseline mutation rate (BMR) based on the intronic mutations in addition to UTRs and flanking intergenic mutations to increase statistical power. This empirical BMR estimate implicitly controls for the heterogeneous mutational landscape across the genome, without the need for inferring BMR from covariate information such as gene expression, replication time and histone marks (see Methods). Further, to control for within-gene variation in mutation risk, DiffInvex employs a locus sampling approach, matching the trinucleotide and pentanucleotide composition to control for mutational signature confounding, and also matches the DNA methylation status, a further determinant of local mutation rates^13^, between target (exonic) and background (intronic and intergenic) regions. Additionally, our DiffInvex implementation filters out problematic genomic regions associated with mutation rate such as low- mappability regions and hypermutated sites (Methods). Finally, we identified and excluded highly recurrently mutated intronic hotspots (Methods) as they are unlikely to have resulted from selection.

**Figure 2.**
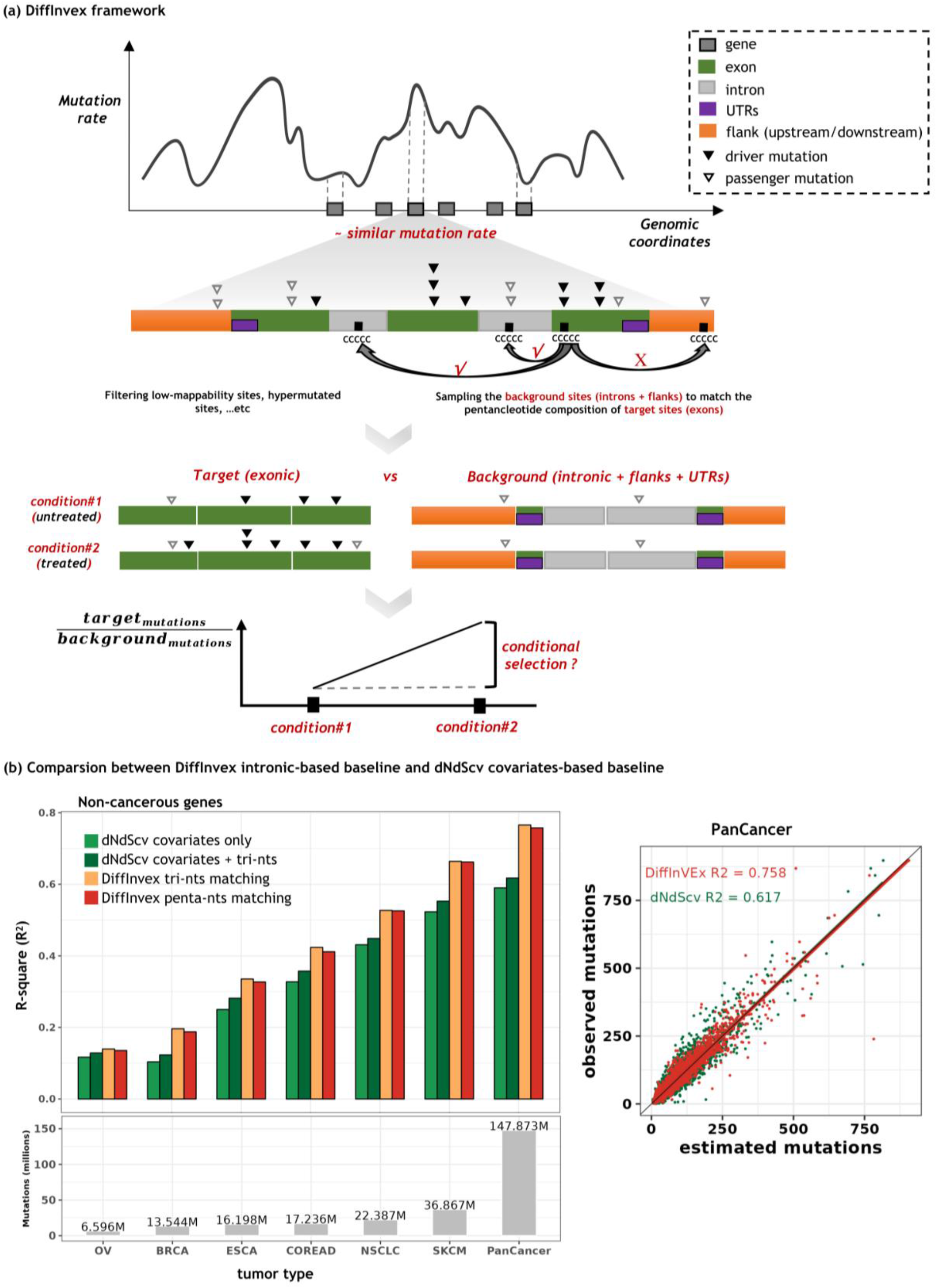
DiffInvex method for quantifying conditional selection in cancer whole genomes. a) A schematic of the DiffInvex framework that utilizes the intronic, UTRs and flanking intergenic regions of each gene as a mutation rate baseline to infer somatic selection and conditional selection. Under no selection, exonic and intronic/UTR regions of the gene have similar mutation rates, after filtering genomic problematic regions and matching their pentanucleotide compositions. DiffInvex normalized the frequency of target (exonic) mutations to background (intronic, UTRs, optionally gene flanks) mutation frequency, comparing two or more conditions to evaluate the conditional selection acting upon that gene. **b)** Accuracy of DiffInvex intronic baseline compared to the baseline inferred from dNdScv covariates in explaining the variation (R^2^) in exonic coding mutation rate for passenger genes in different tumor types. Top-left panel shows the R^2^ of four different models to estimate the exonic mutations: 20 dNdScv covariates only, 20 dNdScv covariates and exonic-trinucleotide compositions, DiffInvex intronic mutation rate after trinucleotide matching between exonic and intronic regions, and DiffInvex intronic mutation rate after pentanucleotide matching between exonic and intronic regions. Bottom-left panel shows the total number of mutations in different tumor types, demonstrating that R^2^ is correlated with the number of mutations. Right panel shows a correlation between the estimated and observed mutation counts in passenger genes (dots) in the pan-cancer cohort, using the DiffInvex with pentanucleotide matching (in red) and the dNdScv with exonic trinucleotide composition (in green).

To evaluate if the DiffInvex empirical BMR could accurately estimate the neutral variation in the mutational landscape between genes, we benchmarked it against modeling of BMR using covariates from dNdScv^14^, a state-of-the-art method for quantifying selection in cancer to identify driver genes. We showed that our DiffInvex empirical BMR achieved the highest R2 in explaining the variation in exonic coding mutation rate for non-cancerous genes, compared to dNdScv covariates in the pan-cancer analysis (DiffInvex using pentanucleotide-matching R^2^ = 0.76, dNdScv covariates R^2^ = 0.62) as well as most tumor types (Fig. 2b top-left panel, Supplementary Fig. 4a, Supplementary Table 3). Across cancer types, the DiffInvex R^2^ as well as the difference between DiffInvex R^2^ and dNdScv R^2^ were positively correlated with the mutation burden (Fig. 2b bottom-left panel), probably reflecting noise resulting from a low number of mutations. This suggests that the benefits of an empirical BMR estimate such as DiffInvex, will grow with the inevitable increase of cancer WGS data. To further support the utility of the intronic BMR estimate, we also applied DiffInvex and the dNdScv tools to the task of identifying cancer drivers in genomes of different tumor types (see Methods). DiffInvex achieved comparable accuracy to dNdScv in detecting known cancer genes in the Cancer Gene Census (CGC) database (Supplementary Fig. 4b). Finally, we evaluated the contribution of DiffInvex modules such as methods for filtering problematic genomic regions, background region width, and use of DNA methylation-based matching, towards accuracy of modeling the neutral BMR (see Methods, Supplementary Fig. 5). We found that DiffInvex filters and DNA methylation-based matching improved the R^2^ by 2.5% and 1%, respectively.

The task of DiffInvex is to determine the differential excess of point mutations in target regions (here, coding exons) over the baseline regions (here, introns and gene flanks) between two conditions or time points (e.g. pre-treated vs treatment-naive, responding vs relapsing). To that end, DiffInvex utilized a Poisson regression model for count data, further regularized by a weakly-informative prior to stabilize estimates from sparse data (Methods). Moreover, the regression model can control for confounding factors between conditions, such as tumor types, which can be differently abundant between pre-treated and treatment-naive samples (Fig. 1a and Supplementary Fig. 1d), and it can control for technical variation between data sourced from different cohorts (Supplementary Fig. 1c). Importantly, the design allows us to control for conjoint drug treatments wherein one drug treatment confounds gene association testing with the other frequently co-administered drugs.

### Identifying chemotherapy-associated drivers via DiffInvex analysis of cancer genomes

We utilized DiffInvex on a large-scale dataset of 8,591 cancer WGS and 147.87 million somatic mutations to identify putative drug resistance genes (DRGs) associated with each drug type, genes that gain positive selection through treatment; additionally, putative drug sensitivity genes (DSGs) that lose positive selection are identified. First, in a pan-cancer analysis, we applied DiffInvex to call these drug type-specific putative DRGs by comparing the mutation profiles of patients exposed to a specific group of drugs (e.g. platinum-based drugs) and treatment-naive patients, while we control for confounders including tumor type, cohort, tumor stage (primary or metastatic) and patient sex. For each gene, 15 DiffInvex regression tests were performed to assess the interaction with each drug type separately.

As a positive control, we asked whether DiffInvex can identify previously-reported genes in which occurrence of point mutations can grant resistance to targeted therapy drugs. Indeed, we identified association of mutations in the *ESR1* gene with prior exposure to antiestrogen drugs, mutations in *AR* gene with antiandrogen drugs, *EGFR* gene with EGFRi drugs, and *KIT* with kinase inhibitor drugs (all effect sizes > 2, all FDRs < 1e^-8^; the “effect size” here refers to natural log fold-enrichment of coding exonic mutation rates in pre-treated samples compared to treatment-naive, normalizing for the change in intronic mutation burdens) (Supplementary Fig. 6a). However, in the initial analysis that considered every drug type separately, we observed that some putative DRGs were strongly associated with many drug types (Supplementary Fig. 6b), which is implausible. For example, *ESR1* gene mutations were also strongly associated with 7 drug types including nitrogen mustards and other alkylating, cytotoxic antibodies, topoisomerase inhibitor, kinase inhibitors, microtubule and pyrimidine (all FDR < 1e^-10^), which might be because most BRCA patients were treated with antiestrogen drugs in combination with other drugs (Fig. 1c). Similarly, *APC* and *SMAD4* tumor suppressor genes and *PIK3CA* and *BRAF* oncogenes were significantly (all FDR < 25%) associated with 10, 4, 6, and 3 drug types respectively. These results showed that conjoint drug treatment, with only 548/2730 patients pre-treated with only one drug type, likely confounded this drug-target association study.

To overcome this challenge, we applied DiffInvex regression to quantify conditional selection associated with each drug type and evaluate its contribution to drug resistance by a joint analysis comparing the mutation profiles of patients exposed to different combinations of the 15 drug types (single and multiple drug types) with treatment-naive patients. For each gene, DiffInvex assesses the contribution of each drug type towards the excess of selected mutations upon treatment simultaneously in a single regression test (Methods).

This allowed DiffInvex to control for the confounding introduced by the conjoint drug treatments as well as tumor type and tumor stage (primary or metastatic). The positive control associations remained (Fig. 3a, Supplementary Table 4) including *ESR1* mutations associated with antiestrogen drugs (effect size = 3.1, FDR = 5.77e^-29^), *EGFR* mutations with EGFR inhibitor drugs (effect size = 2.71, FDR = 1.02e^-25^), *KIT* mutations with kinase inhibitor drugs (effect size = 1.99, FDR = 1.5e^-9^) and *AR* mutations with antiandrogen drugs (effect size = 1.83, FDR = 3.58e^-7^). In the revised analysis, we do not anymore see *ESR1*, *APC* and *SMAD4* mutations significantly associating with multiple drug types (1, 2 and 1 associations at FDR < 25% respectively), suggesting that the joint drug treatments are not strongly confounding. Further, a quantile-quantile plot of the p-values from the association analysis suggested that the p-values were conservative (inflation factor lambda=0.54, Supplementary Fig. 6c), meaning that false positive rates are stringently controlled, albeit at the expense of some false-negatives i.e. missed associations.

**Figure 3.**
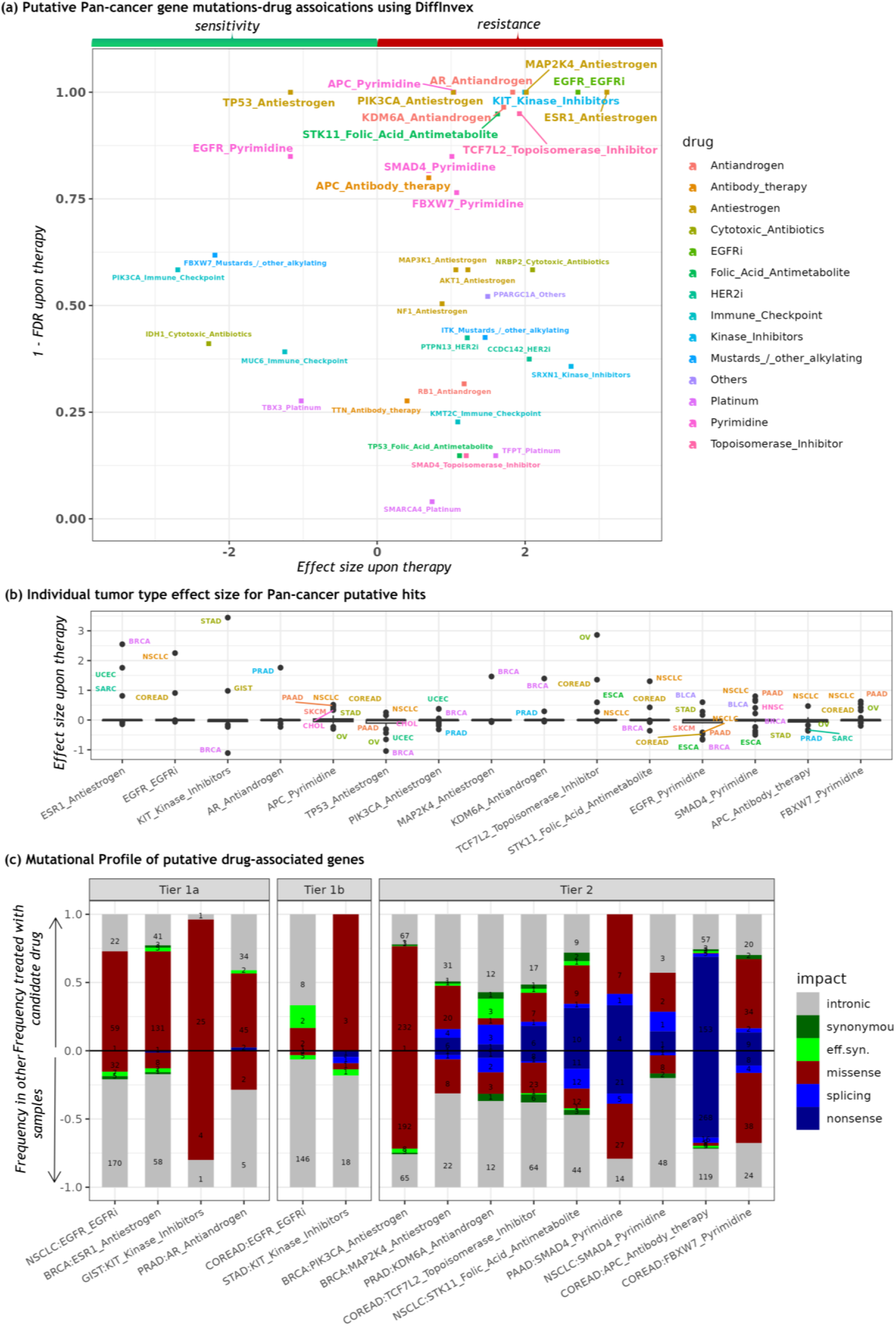
Chemotherapy-associated mutational driver genes identified by DiffInvex. a) Drug sensitivity and resistance associations identified in the pan-cancer analysis of 8,591 cancer genomes, while controlling for confounding between joint drug treatment. At FDR < 25%, there are 13 drug resistance and 2 drug sensitivity associations. Associations are colored by the drug type. The x-axis represents the effect size upon treatment: the natural log fold-enrichment of coding exonic mutation rates in tumor samples pre-treated with the putative drug type compared to other samples, normalizing for the difference in intronic mutation burdens between two groups of samples. **b)** Effect sizes of associations with treatment, derived from the DiffInvex analysis in individual tumor types (listed with acronyms next to corresponding points), here shown for the significant pan-cancer associations. **c)** Mutation functional impact profile of putative drug- associated genes in candidate tumor types. The upper-half of the plot (positive values) represents the fractions of different mutation types (nonsense, splicing, high-impact missense, effectively synonymous (low-impact missense), synonymous, and intronic) in the candidate gene in tumor samples treated with a certain the putative drug type, as listed on the x-axis. The lower-half of the box plot (negative values) represents the fractions of these mutation types in that gene in other tumor samples (treatment-naive samples and samples pre-treated with other drug types). The effectively synonymous (“eff.syn.”) mutations are missense mutations with AlphaMissense pathogenicity score < 0.39. Tier 1 encompasses known drug-gene mutation associations, where Tier 1a does so in known tumor types while Tier 1b includes additional tumor types. Tier 2 has candidate new gene-drug associations with DiffInvex FDR < 25% in the pan-cancer analysis. Please see Supplementary Fig. 6 for similar plots for the more tentative Tier 3a and 3b associations.

In additional to these known positive control genes, DiffInvex identified additional treatment-associated genes with different drug types (11 gene-drug associations with FDR < 25%, of which 9 with resistance and 2 with sensitivity; Fig. 3a). Two general properties of this tail of drug resistance genes stand out. First, they are all known oncogenes and tumor suppressor genes, meaning they are under positive selection even without therapy exposure. (Of note, our methodology is in principle able to identify genes which are selected only after therapy but are not otherwise mutational cancer drivers, such as the *AR* gene. Indeed, the medium-confidence tier of hits with FDR=25-80% does contain 2 such genes, *NRBP2* and *PPARGC1A*). Second, their conditional selection effect sizes are lower (natural log mutation rate enrichments all <=2) than those of the two common drug resistance mutations, the *ESR1* in relation to antiestrogen drugs and *EGFR* mutations in relation to EGFRi drugs (∼2.5).

The specific associations we identified at FDR<25% include those of mutations in *PIK3CA* and *MAP2K4* with resistance to antiestrogen drugs, *APC*, *SMAD4* and *FBXW7* mutations with resistance to pyrimidine drugs, *TCF7L2* mutations with resistance to topoisomerase inhibitor drugs, *KDM6A* mutations with resistance to antiandrogen drugs, and *STK11* mutations with resistant to folic acid antimetabolites. Overall, this suggests that resistance to various anticancer drugs in use today is commonly mediated through driver mutations in known cancer genes, rather than through mutations in specific drug resistance genes, and that they have moderate effect sizes.

### Classification of the identified differentially selected genes by cancer type and degree of prior knowledge

Next, we asked in which individual tumor type these pan-cancer drug-target associations were most relevant, by performing the association testing in individual cancer types and monitoring the effect sizes (Fig. 3b). Based on these results, we divided the observed gene-drug-cancer type combinations into 5 tiers (Fig. 3c and Supplementary Fig. 6d).

First, Tier 1a includes known drug-gene mutation associations in known tumor types such as *ESR1* mutations-antiestrogen resistance in BRCA^15^, *EGFR* mutation-EGFRi resistance in NSCLC^16^, *KIT* mutations-kinase inhibitor resistance in GIST^17^, and *AR* mutations-antiandrogen resistance in PRAD cancer type^18^. These well-known associations are recapitulated in our analyses where they serve as a positive control.

Second, Tier 1b contains known gene mutation-drug resistance (or sensitivity) pairs but includes new tumor types. In other words, these are bona fide drug resistance drivers, however their tissue spectrum is redefined, such that additional tumor types could reap advantages of understanding the drug resistance mechanisms and, possibly, countering them. For example, we find that the *EGFR* mutation - treatment by EGFRi association, which is well-known in NSCLC (effect size [natural log ratio of mutation rates] = 2.25), is relevant to COREAD as well at a suggestive significance threshold (effect size = 0.91). Similarly, our analysis indicates that the *KIT* mutations associated with kinase inhibitors are relevant for STAD (effect size = 3.44) as a Tier 1b hit in our analysis, beside the known Tier 1a relevance to drug resistance in GIST (effect size = 0.98).

Third, Tier 2 has candidate new gene-drug associations with solid statistical support (here, pan-cancer FDR < 25%; we note again that our p-values and therefore also FDRs are likely conservative, see above); these can be associated to certain cancer types by considering the distribution of effect sizes across individual cancer types (Fig. 3b). The Tier 2 encompasses the following observed hits: *PIK3CA* mutations-antiestrogen resistance and *MAP2K4* mutations- antiestrogen resistance, both in BRCA; *KDM6A* mutations-androgen deprivation resistance in PRAD, *TCF7L2* mutation-topoisomerase inhibitor and *APC* mutation-antibody therapy resistance, both in COREAD; *STK11* mutations-folic acid antimetabolites resistance in NSCLC; *SMAD4* mutations-pyrimidine analog resistance in PAAD. Some of these mutation-drug associations were supported by a statistical test on the functional impact of mutations as well as replicated in independent cohorts of pre-treated tumors (see below).

Fourth, Tier 3 includes additional putative gene mutation-drug associations, but with limited evidence of the relevance of those gene mutations in drug resistance (or sensitivity) in individual tumor types. This divides into two categories, Tier 3a that includes putative associations of a frequently-mutated cancer driver gene (pan-cancer FDR < 25%, Fig. 3), however these associations indicate different responses (sensitivity or resistance) in different tumor types (Fig. 3b). For example, *TP53* mutations indicated sensitivity to antiestrogen drugs in the pan-cancer analysis (Fig. 3a, left side), but the *TP53* mutations showed sensitivity and resistance to antiestrogen drugs in BRCA and NSCLC tumors, respectively (Supplementary Fig. 6d). Tier 3a also includes associations of *APC* mutations-pyrimidine analog resistance, and *EGFR* mutations- pyrimidine analog sensitivity in NSCLC and COREAD tumors. Tier 3b includes tentative new gene mutation-drug associations, defined as those hits with FDR<=80% in our main pan-cancer analysis (Fig. 3a, Supplementary Fig. 6d), which could be further validated using the functional impact test and/or in independent panel and whole-exome sequencing cohorts (*MAP3K1*- antiestrogen in BRCA, and *KMT2C*-immune checkpoint in NSCLC) (see below and Fig. 4). Therefore, these associations in Tiers 3a and 3b should be considered with caution, given their more modest FDRs, which indicates higher false positive rates.

**Fig. 4.**
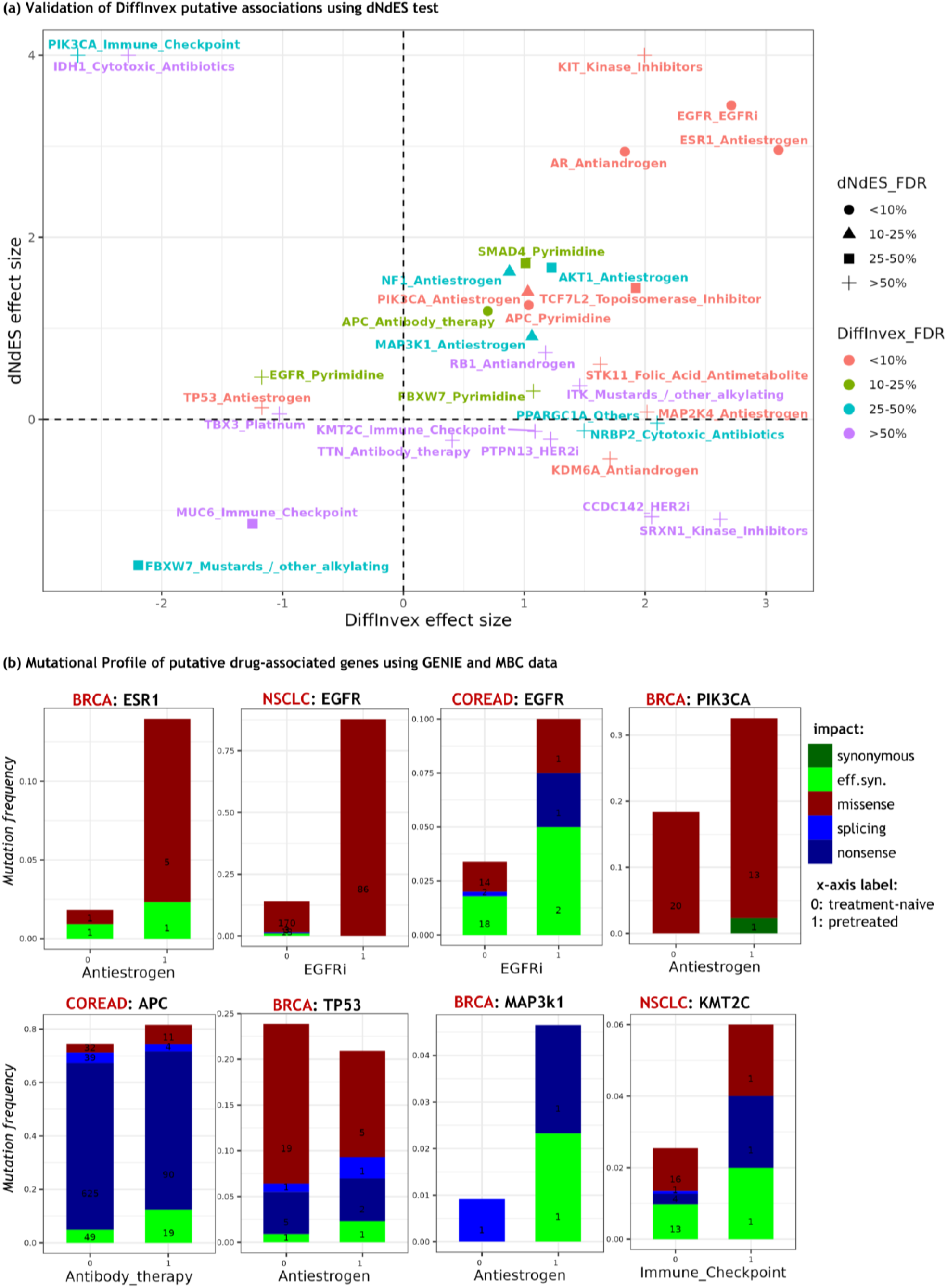
Validation of DiffInvex drug-gene associations in a functional impact test and in independent data sets. a) A test for increased mutation functional impact associated with treatment, dNdES (ES, effectively synonymous) of the pan-cancer associations previously identified by DiffInvex. Scatter plot shows the DiffInvex and dNdES effect sizes of the putative hits. The dNdES effect size is the natural log fold-enrichment of high-impact mutations (nonsense, splicing and high-impact missense i.e. those with AlphaMissense score >= 0.39) in tumor samples pre-treated with the particular drug type compared to other samples, normalizing for the change in low-impact mutations (synonymous and effectively synonymous i.e. low-impact missense). Associations are colored by the FDR range in the pan-cancer analysis by DiffInvex in Fig. 3a. Here, only the DiffInvex gene mutations-drug associations with DiffInvex FDR < 90% were tested using the dNdES test. In this plot, we capped the dNdES selection coefficients at the value of 4. **b)** Replication of some of the DiffInvex significant associations using panel sequencing data from GENIE dataset (for COREAD, NSCLC cancer types) and whole-exome sequencing data from MBC cohort (for BRCA cancer type). In each panel, the title is “tumor type:gene” and the x-axis is the putative drug type. The bar plot shows the mutational impact profile for putative drug- associated genes of the treatment-naive samples (x-axis = 0) and pre-treated samples with that drug (x-axis = 1) in the candidate tumor types. The effectively synonymous (“eff.syn.”) mutations are the low-impact missense mutations that have AlphaMissense pathogenicity score < 0.39.

We further visually inspected the distribution of individual mutations across driver hotspots in the candidate drug resistance genes, contrasting the pre-treated and treatment-naive tumors (lollipop plots in Supplementary Figs. 7-9). In the Tier 1a hits -- known drug-resistance mutations in EGFR, KIT and AR -- do exhibit “specialized” drug-resistance mutations as expected (Supplementary Fig. 7). In contrast, in the Tier 2 and Tier 3 genes the distributions of driver mutations seem broadly consistent across the treated and non-treated tumor genomes (see example of PIK3CA in antiestrogen treated vs untreated BRCA in Supplementary Fig. 8a). These genes and encoded proteins are not the known targets of the drugs we find they are associated with, and appear to confer resistance via an increase in driver gene potential rather than by specialized resistance mutations. While this is the general trend, there might be individual cases that run counter to that (e.g. the Tier 2 gene *MAP2K4* and Tier 3b gene *MAP3K1* with antiestrogen resistance). In case of *MAP2K4* gene, exon 5 seems to be enriched in splice-site mutations in pre-treated tumors, compared to treatment-naive tumors, which could disrupt RNA splicing and protein-coding sequence (Supplementary Fig. 9a). Additionally, in case of *MAP3K1,* exon 10 versus exon 14 appear differentially affected by nonsense mutations, thus generating truncations of different lengths (Supplementary Fig. 9b). However, larger cohorts are needed to test significance.

To further filter out false positives in these Tiers 3a and 3b, and also to validate the more confident Tiers 1a, 1b and 2 above, we next devised and implemented a complementary statistical test for differential selection associated with drug therapy. This test is based on the rationale that if the mutation rate increases after therapy is biologically meaningful, the functional impact of the mutations on the protein sequence should also increase in contrast to untreated.

### Validation of gene mutation-drug treatment associations using dNdES test for differential functional impact

We next implemented a functional impact test named dNdES to confirm the DiffInvex hits, however using a different baseline which does not rely on local mutation rates as estimated from introns and gene flanks. For each gene, the dNdES test compares the frequency of nonsynonymous to “effectively synonymous” (ES) mutations under different conditions, here implying, again, tumor samples treated with a putative drug type versus other tumor samples (treatment-naive samples and samples pre-treated with other drug types). We annotated the exonic mutations with scores from AlphaMissense, a state-of-the-art predictor for missense mutation pathogenicity^19^, thus classifying the missense mutations with score < 0.39 as ES and grouping them with synonymous mutations to form a baseline for estimating functional impact (see Methods). Expectedly, the positive side of the log nonsynonymous to effectively-synonymous ratio (log dNdES) was correlated to exonic-to-intronic ratio (log dEXdIN; Supplementary Fig. 10a), considering general selection on known cancer genes. This correlation remains observed for different cutoff values of the AlphaMissense score applied, where the cutoff of 0.39 that we chose results in a slope ∼=1 for this correlation in somatic mutation rates on cancer genes (Supplementary Fig. 10c). Of note, dNdES ratios have a higher spread compared to DiffInvex ratios (Supplementary Fig. 10b), likely because of more sparse mutation occurrence in the ES loci used as a neutral baseline, compared to the more abundant intronic loci in the default DiffInvex (dEXdIN) approach.

Next, we applied the AlphaMissense-based dNdES test to our task of identifying conditionally- selected genes, comparing the functional impact of exonic mutations in pre-treated tumors relative to treatment-naive tumors. Encouragingly, the effect sizes of conditional selection between the 2 tests correlate (Fig. 4a, Supplementary Table 4), and many DiffInvex putative associations have been validated in the dNdES test (Fig. 4a). Out of 15 DiffInvex significant associations (FDR < 25%), 6 associations were also statistically significant using dNdES test (FDR <25%), 2 were dNdES-significant at a permissive FDR 50%, and additional 4 associations had a consistent direction of response (sensitivity or resistance) as they did in the discovery DiffInvex.

In particular, the known positive control, Tier 1 hits including *ESR1*-antiestrogen, *EGFR*-EGFR inhibitors and *AR*-antiandrogen had strong effect sizes and significance in the dNdES test (Fig. 4a). Additionally, the pan-cancer study of differential dNdES validated some Tier 2 and Tier 3a associations (FDR < 25%) including *APC*-pyrimidine analogs, *PIK3CA*-antiestrogens, and *APC*-antibody therapy drugs, and at a permissive dNdES FDR < 50% additionally supported *SMAD4*- pyrimidine analogs and *TCF7L2*-topoisomerase inhibitors. However, other associations did not reach significance in the (less-powered) dNdES test including associations of *KIT*-kinase inhibitors, *STK11*-folic acid antimetabolites, *MAP2K4*-antiestrogen (marginally), and *FBXW7*- pyrimidine analogs. In some cases, this might be attributed to the low numbers of synonymous and effectively synonymous mutations that are used as a baseline. For example, in testing the known *KIT* mutations-kinase inhibitors association, no synonymous or effectively synonymous mutations were observed in the *KIT* gene in the samples pre-treated with kinase inhibitor drugs (Fig. 3c) and so this association could not be validated but was not invalidated either.

Additionally, in Tier 3b hits, two associations were significant in the dNdES test at FDR < 25% (*NF1* and *MAP3K1*-antiestrogen resistance), one association at dNdES at FDR < 50% (*AKT1* mutations - antiestrogen resistance). Other Tier 3b associations did exhibit the correct direction in the dNdES test, such as *RB1*-antiandrogen resistance, and the *ITK* gene mutations associated with resistance to mustards & other alkylating agents (this drug group excludes platinum). In contrast, some Tier 3a and Tier 3b candidates would be invalidated by the dNdES test, in particular the tentative resistance mutations in *SRXN1* and *CCDC142*, and the tentative sensitizing mutations in *EGFR*-pyrimidine analogs, *PIK3CA*-immune checkpoint drugs and *IDH1*- cytotoxic inhibitors are invalidated by dNdES (Fig. 4a). As a caveat, the dNdES test relies on AlphaMissense and may not be informative in case of genes or sites where AlphaMissense does not perform well; these could be some of the cases of genes with ∼0 dNdES effect size (Fig. 4a). Several genes show this effect size such as *TBX3* with platinum sensitivity, *KMT2C* with immune checkpoint resistance, or *NBRP2* and cytotoxic antibiotics resistance.

The above results provided support for the DiffInvex approach overall, and supported many individual putative hits identified by DiffInvex by detecting an increase in impactful mutations in pre-treated tumors.

### Validation of gene-drug associations using independent cohorts

Next, we sought to replicate the DiffInvex putative DRG associations using panel and whole-exome sequencing data from independent studies with available treatment information. This includes the NSCLC and CRC cohorts from the GENIE Biopharma Collaborative (BPC) project^20^, as well as the BRCA cohort from the metastatic breast cancer genomics project^21^. Data from other cancer types (including gene sequencing of pre-treated tumors and treatment metadata) was to our knowledge not available to us at the time of writing, so other cancer types apart from breast,

colon and lung were not tested in the replication. We obtained the mutation profiles of 1747, 1469 and 186 samples from NSCLC, COREAD and BRCA cohorts, respectively, which were either treatment-naive or pre-treated (Methods). For each putative gene-drug association we identified in the discovery cohort, we compared in the replication cohort the frequency of mutations in this gene in the pre-treated samples with that drug versus the treatment-naive samples.

Our analysis considered only the coding mutations and splice site mutations, since these were not WGS datasets. Moreover, we stratified the missense mutations into low-impact (effectively- synonymous, ES) and high-impact missense based on AlphaMissense scores. Next, we compared proportions of different mutation impact categories as in the dNdES test as above.

This analysis confirmed some associations for different tumor types (Fig. 4b), most prominently in the external breast cancer dataset^21^, henceforth the MBC dataset. This included the positive control *ESR1*-antiestrogen associations, as well as the discovered associations herein *PIK3CA*- antiestrogen, *TP53*-antiestrogen and *MAP3K1*-antiestrogen. For example, in case of *PIK3CA*- antiestrogen resistance, 18% (20 out of 109) treatment-naive tumors have high-impact missense *PIK3CA* mutations, while 30% (13 out of 43) of pre-treated tumors bear high-impact missense mutations. Additionally, in case of *MAP3K1*-antiestrogen treatment resistance, the frequency of high-impact mutations increased from 9% in treatment-naive patients to 23% in the pre-treated samples. The putative *MAP2K4*-antiestrogen association could not be assessed in this data set as we observed no *MAP2K4* mutations in the pre-treated patients with antiestrogen drugs, but please see below for a validation of *MAP2K4* and other breast mutations in longitudinal genomics datasets. As another example from the independent MBC cohort, we highlight *TP53* mutation- antiestrogen treatment sensitivity, which had high DiffInvex significance (albeit not supported in dNdES in the discovery cohort; Fig 4a). In treatment-naive tumors 96.1% of the *TP53* mutations are high-impact (missense, nonsense or splice) while 3.9% are low-impact (effectively synonymous by our criteria), while in pre-treated tumors 88.9% are high and 11.1% are low impact. Thus, the frequency of *TP53* high-impact mutations modestly decreased from 22.3% (25 mutations in 109 samples) in treatment-naive patients to 18.6% (8 mutations in 43 samples) in pre-treated patients with antiestrogen drugs.

Next, for non-small cell lung cancer we considered the GENIE cohort as external validation. Here, the positive control *EGFR*-EGFR inhibitor association was evident, as well as the link between *KMT2C* mutations and immune checkpoint inhibitors (Fig. 4b). In case of *KMT2C*-immune checkpoint treatment resistance, in treatment-naive tumors 61.8% of the *KMT2C* mutations are high-impact (high-AlphaMissense score missense, nonsense or splice), while in pre-treated tumors 66.7% are high impact. The frequency of high-impact mutations increased from 1.6% (21 mutations in 1334 samples) in treatment-naive tumors to 6% (3 mutations in 50 samples) in tumors pre-treated with immune checkpoints drugs. We could not validate the putative *STK11* mutations-folic acid antimetabolites.

In the GENIE colorectal cancer cohort, we note support for our Tier 1b association between *EGFR* mutations and pre-treatment with EGFR inhibitors. In this case, the frequency of high-impact mutations increased from 1.6% (16 mutations in 1001 samples) in treatment-naive tumors to 5% (2 mutations in 40 samples) in pre-treated tumors. Further, the link between *APC* mutations and pretreatment with antibody therapy (commonly, the Bevacizumab drug, 349 out of 398 cases of the antibody therapy in our cohort are bevacizumab) was supported in the GENIE colorectal cohort (bottom-left panel Fig. 4b). Here, the frequency of all mutations increased from 74.4% (745 mutations in 1001 samples) in treatment-naive to 81.6% (124 mutations in 152 samples) in pre-treated tumors, but the frequency of high-impact mutations was 69% in both treatment-naive and pre-treated tumors.

Moreover, results from other smaller-scale studies in breast cancer patients showed support of some DiffInvex associations. Using a longitudinal study of 48 patients on aromatase inhibitors (a type of antiestrogen therapy), Lopez-Knowles et al.^22^ reported that *MAP2K4* mutations were observed in 1 patient post-treatment with antiestrogen drugs, but not pretreatment. Additionally, Brady et al.^23^ showed that out of 3 patients treated with antiestrogens, two patients gained *ESR1* mutations and one patient gained *MAP2K4* mutation upon treatment. Similarly, Huang et al.^24^ suggested that *PIK3CA* mutations contributed to fulvestrant resistance in ER-positive breast cancer as they observed mutations in 3 out of 4 fulvestrant-resistant patients. In another study using ctDNA from 141 advanced breast cancer patients that underwent first-line standard treatment, Liao et al.^25^ showed that *PIK3CA* and *TP53* variants were associated with drug resistance. *PIK3CA* and more generally the PI3K pathway has convergent evidence, including from the clinic, for being causally involved in breast cancer resistance to endocrine therapies (reviewed in Araki et al.^26^ and Rusquec et al.^27^). In a clinical study, the inhibitor of the *PIK3CA*- encoded p110α, alpelisib, potentiated the effects of fulvestrant therapy significantly only in the *PIK3CA*-mutant breast tumors^28^. Analysis of compiled data from three earlier clinical trials indicated that tumors with *PIK3CA* mutations had a lower response rate to antiestrogen therapy^29^. Our large-scale genomics analysis supports that *PIK3CA* mutations preferentially occur after therapy, implicating this gene in antiestrogen resistance. Overall, in breast cancer, we identified associations of *PIK3CA*, *MAP2K4*, and *MAP3K1* mutations with resistance to antiestrogen drugs as well as associations of *TP53* mutations with sensitivity to antiestrogen drugs, which were replicated in the MBC cohort and/or in other smaller-scale studies.

Further smaller scale studies provide support for some identified hits in colon and prostate cancer. For prostate cancer, Grasso et al.^30^ reported that *KDM6A* was altered in pre-treated metastatic castration-resistant prostate cancer (CRPC) with 3 copy number gains, 2 losses, and 1 point mutations in CRPC compared to 0 alterations in treatment-naive prostate cancer. Using cell lines, Das et al.^31^ showed that CRC with *APC* mutations developed resistance to 5-FU (pyrimidine analog by our categorization). Using CRC patients and cell lines, two studies suggested that loss of *SMAD4* in CRC patients induces resistance to 5-FU-based therapy^32,33^, apparently consistent with our association of *SMAD4* selected mutations with pyrimidine analogs pretreatment.

## Discussion

Mechanisms by which drug resistance arises in cancer are important to understand, such that the knowledge of these mechanisms may be applied to prevent (or target) them. Genome sequences of tumors that were pre-treated by chemotherapies will reflect evolutionary pressures to resist the therapy and should therefore contain resistance-conferring mutations. Genome sequences of tumors not pre-treated should either not contain such mutations, or they should contain them but signatures of selection on such mutations would be weaker. Therefore, a rigorous comparison of genomes of tumors before versus after treatment is, in principle, a powerful tool to identify drug resistance mutations^34^. However, this approach is fraught with challenges, in particular the mutation rate heterogeneity and the changes in mutation spectra caused by therapy^9,35^, which mean that changes in mutation burdens of individual genes could result from neutral processes. We rigorously addressed these challenges by the DiffInvex framework, and in addition accounted for confounding between various treatments, suggesting putative drug resistance driver mutations. However, our study has some limitations. One stems from limited statistical power to identify the less common conditionally-selected genes associated with chemotherapy. This will be addressed by increases in the amount of cancer WGS data generated from different projects, such as the Genomics England with currently ∼14 k WGS^36^, however the lack of genomes from pre-treated tumors remains a bottleneck. Additional data will make it possible to conduct our study to test significance on individual tumor types (currently we tested pan-cancer), which may result in discovering more associations with lower-abundance chemoresistance mutations, giving more insight into whether there exists a “long-tail” of drug resistance driver point mutations. Such lower abundance chemoresistance mutations would exist because their effects on tumor fitness under therapy are milder, or maybe in some cases if the effects are strong but they are relevant only to a rare tumor type or subtype. Additionally, our DiffInvex framework implements an orthogonal “dNdES” method that can be used for validation by testing differences in functional impact; this method will, because of rarity of exonic synonymous or effectively-synonymous mutations, benefit even more with additional genome sequences in the future, and it will also benefit from future iterations of mutation effect predictors that supplant AlphaMissense as the field-leading algorithm. A further limitation, which is conceptual in nature and merits development of additional methodologies, is that drug resistance in tumors may in many cases not evolve via point mutations, but instead via copy number alterations and/or epigenetic changes, which DiffInvex is not designed to model. Indeed, our results may reflect the fact that point mutations are not necessarily the dominant mechanism generating drug resistance in tumors. In particular, in addition to the well-known resistance drivers to targeted therapies (*EGFR*, *ESR1*, *AR* and *KIT* mutations), other chemoresistance drivers we identified have lower effect sizes. Moreover, they are largely mutations in known cancer driver genes like PIK3CA, APC and MAP3K1 that associate with drug resistance, rather than mutations in genes particular to drug resistance (which we do not exclude, e.g. *ITK*, *TFPT* and *NRBP2* are tentative hits, which however await replication in larger cohorts; *TFPT* association with platinum resistance seems plausible given its roles in nucleotide excision repair and in promoting apoptosis). Of note, even if mutations in known driver genes are driving therapy resistance, rather than drug resistance mutations in specialized genes or direct drug targets, such mutations with both driver and chemoresistance roles are nonetheless useful for stratifying patients for therapy in a genomically-informed manner. Further applications of our DiffInvex method to ever-growing datasets will increase the repertoire of mutational cancer chemoresistance driver genes, and thus further efforts for personalized medicine.

## Methods

### Data collection and processing

We collected WGS somatic mutation data from nine different studies (Supplementary Table 1). First, we obtained somatic SNVs for 4,858 WGS from the Hartwig Medical Foundation study (https://www.hartwigmedicalfoundation.nl/en/)^3^. Second, we downloaded somatic SNVs from 2,808 WGS from the Pan-cancer Analysis of Whole Genomes (PCAWG) study that were re- processed by the Hartwig pipeline at the International Cancer Genome Consortium (ICGC) data portal (https://dcc.icgc.org/releases/PCAWG/Hartwig)^34^. Third, we downloaded somatic SNVs from 570 WGS from the Personal Oncogenomics (POG) project from BC Cancer (https://www.bcgsc.ca/downloads/POG570/)^5^. Fourth, we obtained somatic SNVs for 724 WGS from the DECIDER study (https://www.deciderproject.eu/)^6^. Fifth, we downloaded somatic SNVs for 133 ovarian WGS from the British Columbia Ovarian Cancer Research Program (herein referred to as OVCARE)^37^. Sixth, we downloaded somatic SNVs from the breast and prostate projects that were not included in the PCAWG study from the ICGC data portal (https://dcc.icgc.org/releases/release_25/Projects). These ICGC samples included 352 samples from BRCA-EU project^38^, 183 samples from PRAD-CA project, 91 samples from PRAD-UK project, and 152 samples from EOPC-DE project. We also downloaded read alignments (BAM files) of WGS samples for 907 tumors from the MMRF COMMPASS project^39^ and for 548 tumors from the Clinical Proteomic Tumor Analysis Consortium 3 (CPTAC-3) project^40^ from the GDC data portal (https://portal.gdc.cancer.gov/). Finally, we downloaded the BAM files for 610 samples from the Mutographs project^41^ from the European Genome-Phenome archive (https://ega-archive.org/datasets/EGAD00001006732). Somatic variants of MMRF-COMMPASS, CPTAC-3 and Mutographs projects were called using Strelka2 caller with SomaticEVS score ≥6^42^. Then, weperformed a liftOver of mutation calls from the GRCh38 to the hg19 reference genomes for those samples, as well as samples from the DECIDER project.

We filtered these tumor WGS to exclude tumor samples with missing treatment or tumor stage information. Then, we also excluded samples with tumor purity < 20%, and ultramutated samples from microsatellite instability (MSI) tumors and colorectal tumors with POLE mutations, and finally also the problematic Hartwig and PCAWG samples that were reported in earlier studies using the same data^34,43^. For the PCAWG cohort, we excluded samples tagged with PCAWG blacklist, No pipeline output, Corrupt HMF pipeline 5 run, No PURPLE 2.53 or LINX 1.14 output, and SV INV outlier. In the Hartwig cohort, we excluded one sample with < 50 SNVs and 108 samples from unknown primary sites. Additionally, we filtered out one duplicate PCAWG patient (DO217844) that was also included in the Hartwig cohort. In the MUTOGRAPH cohort, we additionally filtered out 7 ESCA and 2 COREAD ultramutated samples with > 1 million SNVs. From the CPTAC3 cohort, we kept 336 samples from NSCLC, PAAD, RCC, and UCEC (out of 436 samples with purity >= 20%). Finally, we selected one sample or one time point per patient for DECIDER and Hartwig patients with multiple tumor samples. In case of DECIDER cohort, we kept one time point per patient by merging samples from different sites, removing redundant mutations, at the most recent biopsy time point, resulting in 202 genomes from DECIDER. In the Hartwig cohort, if available, we kept one pre-treated sample per patient.

For the validation datasets, we downloaded mutation calls for 3934 samples across 3 cancer types. We obtained somatic SNVs of panel sequencing data from 2004 COREAD samples (https://www.synapse.org/#!Synapse:syn30991602) and 1551 NSCLC samples (https://www.synapse.org/#!Synapse:syn27056179) from the GENIE Biopharma Collaborative (BPC) project (https://genie.cbioportal.org/)^7^. Additionally, we downloaded somatic SNVs of whole-exome sequencing data from 379 BRCA samples from the metastatic breast cancer genomics project (https://www.cbioportal.org/study/clinicalData?id=brca_mbcproject_2022)^21^. In the GENIE cohort, similarly we kept one sample per patient, at the most recent biopsy time point, resulting in 1747 NSCLC samples (413 pre-treated and 1334 treatment-naive) and 1469 COREAD samples (468 pre-treated and 1001 treatment-naive). For the BRCA longitudinal cohort, we excluded blood samples and tissue samples with missing treatment information. For patients with more than one sample, we kept only the most recent sample. This resulted in 109 treatment-naive and 77 pre-treated BRCA samples in the validation dataset.

### Baseline for neutral mutation rate estimation

To estimate the neutral mutation rate in protein-coding sequence (CDS) regions, we utilized the mutations in their neighboring regions including introns, untranslated regions (UTRs), and upstream and downstream gene flanking regions. We utilized the Ensembl annotation package (EnsDb.Hsapiens.v75) to get the coordinates of genomic regions (CDSs, exons, introns, and UTRs) in the hg19 reference genome. Starting with the HGNC symbol of the gene, we selected the most-expressed transcript^44^ if available. Otherwise, we chose the longest transcript. We then defined the target regions (regions of interest) as the CDSs and the adjacent five nucleotides of each intron (to account for splice site disrupting mutations), while the background regions included intronic, UTRs, and intergenic regions, but excluding the coding exonic regions of neighbor genes.

In the case when the total intronic regions were shorter than the minimum required background region size (user-defined parameter, default value = 50,000 nucleotides), we included adjacent genomic loci from the upstream and downstream intergenic regions into the background.

Then, we implemented many filters to exclude genomic loci that could confound the estimation of neutral mutation rate. This includes microsatellite repeats of 6 or more nucleotides, CTCF and cohesin (RAD21) overlapping binding sites, the ENCODE blacklist of problematic regions of the genome^45^, non-uniquely mappable sites according to the Umap k100 alignability track (https://bismap.hoffmanlab.org/)^46^, target regions of Activation-induced cytidine deaminase (AID) somatic hypermutation process (see Supek et. al.^47^ and references therein), and promoter hypermutated sites (NYTTCCG motifs within promoters)^48^. Finally, we filtered out the loci in the background regions bearing hotspots that have 3 or more mutations in our cohort, suggestive of locally high mutation risk.

Next, to minimize the variation in mutation risk due to mutational signatures differing between the target regions and background regions, we employed a locus sampling approach to match their trinucleotide or pentanucleotide (the default) compositions as well as DNA methylation status, within each gene. We iteratively removed nucleotide positions from the background regions to increase similarity in composition to the target regions, until reaching a tolerance < 0.01 (Euclidean distance between the relative trinucleotide/pentanucleotide frequencies in the DNA sequence composition of each region).

### The DiffInvex test for somatic selection and conditional selection

After defining the target and background regions for each gene, we utilized Poisson regression to model the mutation counts in each region and to determine the effect estimates of conditional selection using the following model:

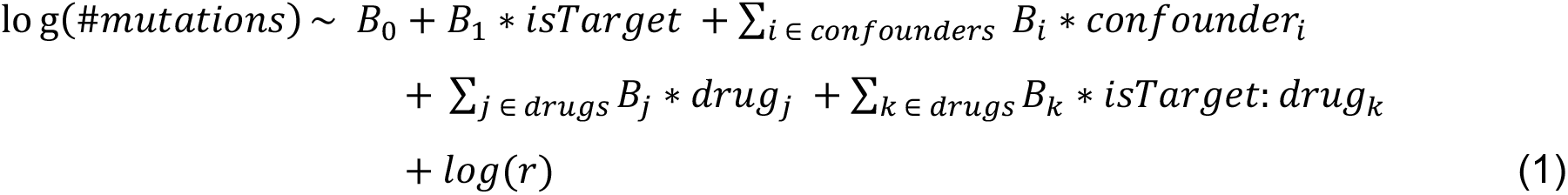

where regression coefficient 𝐵_0_ represents the base mutation rate and is included as the intercept of the model. The 𝑖𝑠𝑇𝑎𝑟𝑔𝑒𝑡 is a binary indicator variable to distinguish mutations in target regions (𝑖𝑠𝑇𝑎𝑟𝑔𝑒𝑡 = 1) that are being tested for selection and mutations in the background regions (𝑖𝑠𝑇𝑎𝑟𝑔𝑒𝑡 = 0) that represent the neutrally accumulated mutations. The corresponding regression coefficient 𝐵_1_ represents the natural log fold change of mutation rates in the target regions (CDS and splicing sites), compared to mutation rates in the background regions (introns, UTRs and, optionally, intergenic regions). Therefore, for each gene, positive 𝐵_1_ indicates that gene is under positive selection in cancer as there is enrichment of mutations in target regions compared to the baseline (background regions), while negative 𝐵_1_ suggests that it is under negative selection in cancer as there is mutation depletion in target regions. The differences in trinucleotide or pentanucleotide compositions between the two regions are controlled by locus sampling to remove parts of the background region (see above).

The 𝑐𝑜𝑛𝑓𝑜𝑢𝑛𝑑𝑒𝑟_𝑖_ represent the variables that could confound our analyses and should be controlled for, and the regression coefficients 𝐵_𝑖_ reflects their effects (for binary variables, the fold-difference in mutation risk between the 2 levels of the variable, on the natural log scale). In our analyses, we adjusted for several confounders including tumor type in the pan-cancer study, the source study, the tumor stage (primary or metastatic), and the patient sex. Similarly, regression variable 𝑑𝑟𝑢𝑔_𝑗_ represents the treatments that were given to different patients in our cohort (𝑑𝑟𝑢𝑔_𝑗_= 1 meaning this drug *j* was given to patients prior to treatment, and 0 if not given prior to therapy) and the coefficient 𝐵_𝑗_ reflects their effects on the background mutation rate (on the natural log scale).

The terms 𝑖𝑠𝑇𝑎𝑟𝑔𝑒𝑡: 𝑑𝑟𝑢𝑔_𝑘_ represent the interaction terms of the target-region indicator variable 𝑖𝑠𝑇𝑎𝑟𝑔𝑒𝑡 with the given treatment indicator variable (𝑑𝑟𝑢𝑔_𝑘_); the corresponding regression coefficients 𝐵_𝑘_ reflects the conditional selection effects by estimating the difference of the natural log fold change of target region to background region mutation rates between the two conditions of the 𝑑𝑟𝑢𝑔_𝑘_ (0: drug treatment was absent, 1: drug treatment was given). So, for 𝑑𝑟𝑢𝑔_𝑘_, positive 𝐵_𝑘_ indicates that the tested gene is a putative drug-resistance gene, as there is an increase of log ratio of target-to-background region mutations when patients that were pre-treated with that drug (𝑑𝑟𝑢𝑔_𝑘_= 1) compared to patients untreated with that drug (𝑑𝑟𝑢𝑔_𝑘_= 0). In contrast, negative 𝐵_𝑘_ indicates that the tested gene is a putative drug-sensitivity gene as there is a decrease of log ratio of target-to-background region mutations when patients were pre-treated with that drug (𝑑𝑟𝑢𝑔_𝑘_= 1) compared to patients untreated with that drug (𝑑𝑟𝑢𝑔_𝑘_= 0). Finally, we include the number of nucleotides-at-risk 𝑟 as an offset in the regression model to normalize mutation counts to relative mutation rates (per nucleotide per sample). This allows DiffInvex to account for variations in DNA lengths of the target and the background regions (after tri/pentanucleotide sampling), as well as the number of samples of each condition (e.g. number of primary versus metastatic samples, or number of samples pre-treated by one drug versus untreated ones). We implemented the regression model as a generalized linear model with a log link function, further regularizing the regression coefficients by using a weakly informative prior to stabilize estimates from sparse data^49^ (using the R function bayesglm, arm package version 1.11_2), as applied in Besedina et al.^50^. The statistical test applied on the regression coefficients (log enrichments) was the Z-test, two-tailed, as implemented in the R function summary(). Multiple testing correction was performed using the FDR method separately for the selection coefficients (𝐵_1_) and for the conditional selection coefficients (𝐵_𝑘_).

In our analyses, we applied the DiffInvex model in two scenarios to obtain the conditional selection effects of each gene-drug interaction. In the first scenario, we tested the associations between gene mutations and each drug type individually, without controlling for the conjoint drug treatment problem:

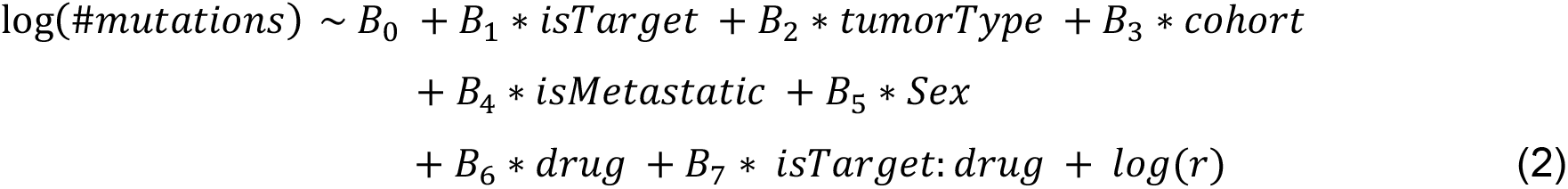

where 𝑑𝑟𝑢𝑔 represents the drug type to be tested (e.g. platinum drugs) and 𝐵_7_ reflects its conditional selection effect (in the natural log scale). Here, for each gene, we applied the DiffInvex mode 15 times (one for each drug type). Results for this scenario were shown in Supplementary figure 6a, and they likely result in many spurious associations.

Therefore, we devised the second testing scenario, where we tested the associations between gene mutations and all drug types in the same model to control for the conjoint drug treatment problem:

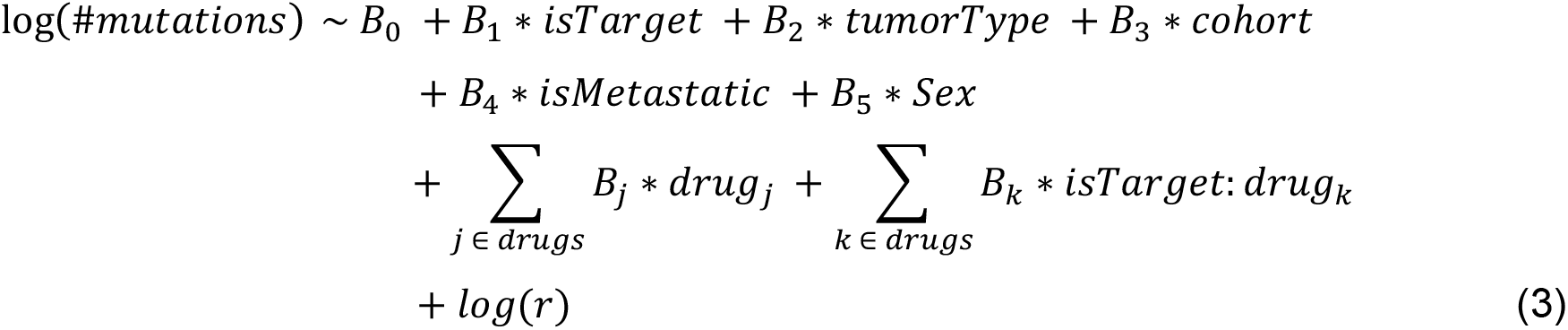

where 𝐵_𝑘_ reflects its conditional selection effect (in the natural log scale) of the drug type 𝑑𝑟𝑢𝑔_𝑘_. We applied the DiffInvex model once for each gene, and results for this scenario were shown in Figure 3a for the pan-cancer analysis, and Figure 3b shows the coefficients for the selected associations in the cancer type-specific analyses.

### The dNdES statistical test for conditional mutation impact on coding regions

We also implemented the dNdES test, to test if a putative DiffInvex gene (DRG or DSG) is under conditional selection upon drug treatment, using a different test on the mutations that does not rely on the intronic baseline, and therefore provides a validation method for DiffInvex. The dNdES functional impact test compares the frequency of nonsynonymous “N” (high-impact missense, nonsense, splicing) to effectively synonymous “ES” (low-impact missense and synonymous) exonic mutations. For that, we first annotated the exonic mutations using Annovar tool^51^ and AlphaMissense method^19^. The AlphaMissense provides a score for missense mutation pathogenicity. We defined the AlphaMissense score = 0.39 as the threshold for dividing missense mutations into high-impact and low-impact missense mutations. This AlphaMissense score resulted in approximately unity slope in the linear regression model between dEXdIN (exonic to intronic mutations) and dNdES (nonsynonymous to effectively synonymous mutations) of cancer genes as shown in Supplementary Fig. 10c. We used this threshold for annotating the missense mutations from the WGS data (discovery cohort) and from the panel and whole-exome sequencing data (validation cohort).

We again utilized Poisson regression to model the mutation counts in exonic regions (target, background) and to determine the effect estimates of selection and conditional selection, for each drug separately, using the following model:

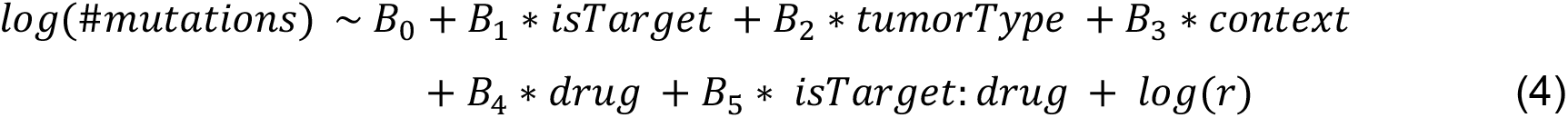

where regression coefficient 𝐵_0_ represents the base mutation rate and is included as the intercept of the model. Regression coefficient 𝑖𝑠𝑇𝑎𝑟𝑔𝑒𝑡 is a target variable to distinguish mutations in target regions (𝑖𝑠𝑇𝑎𝑟𝑔𝑒𝑡 = 1) that is currently being tested for selection and mutations in the background regions (𝑖𝑠𝑇𝑎𝑟𝑔𝑒𝑡 = 0) that represent the neutrally accumulated mutations. In this test, we further classified the exonic sites (importantly, not the mutations), into target regions (sites with 2 or 3 of their possible mutations are nonsynonymous with AlphaMissense >= 0.39) and background regions (sites with 0 or 1 of their possible mutations are nonsynonymous with AlphaMissense >= 0.39). The regression coefficient 𝐵_1_ represents the natural log fold difference of mutation rates in the target regions, compared to mutation rates in the background regions.

The 𝑡𝑢𝑚𝑜𝑇𝑦𝑝𝑒 variable represents the tumor type to control for in the pan-cancer study and 𝐵_2_ reflects its effect (natural log fold difference, relative to one arbitrarily chosen tumor type). Similarly, the 𝑐𝑜𝑛𝑡𝑒𝑥𝑡 variable represents the trinucleotide context (e.g. CCA, CCC, CCG, CCT, ….) of mutations that could confound our analyses and 𝐵_3_ reflects its effect (natural log fold difference, relative to one arbitrarily chosen trinucleotide). The variable 𝑑𝑟𝑢𝑔, the interaction term 𝑖𝑠𝑇𝑎𝑟𝑔𝑒𝑡: 𝑑𝑟𝑢𝑔 and the offset 𝑟 are explained in the DiffInvex test above. We implemented the regression model as a generalized linear model with a log link function (using the R function glm, stats package).

## Acknowledgements

Work in the lab of F.S. was supported by an European Research Council ERC CoG ‘STRUCTOMATIC’ (101088342), European Commission Horizon2020 project ‘DECIDER’ (965193) and Horizon Europe project ‘LUCIA’ (101096473), the Spanish government project ‘REPAIRSCAPE’ (PID2020-118795GB-I00), CaixaResearch project ‘POTENT-IMMUNO’ (HR22-00402), an ICREA professorship, EMBO YIP funding, the SGR funding of the Catalan government (SGR 00616), and a Novo Nordisk Fonden starting grant. We thank Elizaveta Besedina for assistance with developing the DiffInvex Poisson regression model, and Daniel Naro for retrieving and processing (alignment and variant calling) the WGS data from the CPTAC-3, MMRF-COMMPASS, OVCARE and Mutographs studies.

