## Supplementary material for "The DiffInvex evolutionary model for conditional somatic selection identifies chemotherapy resistance genes in 10,000 cancer genomes": Supplemenary Figures 1-10

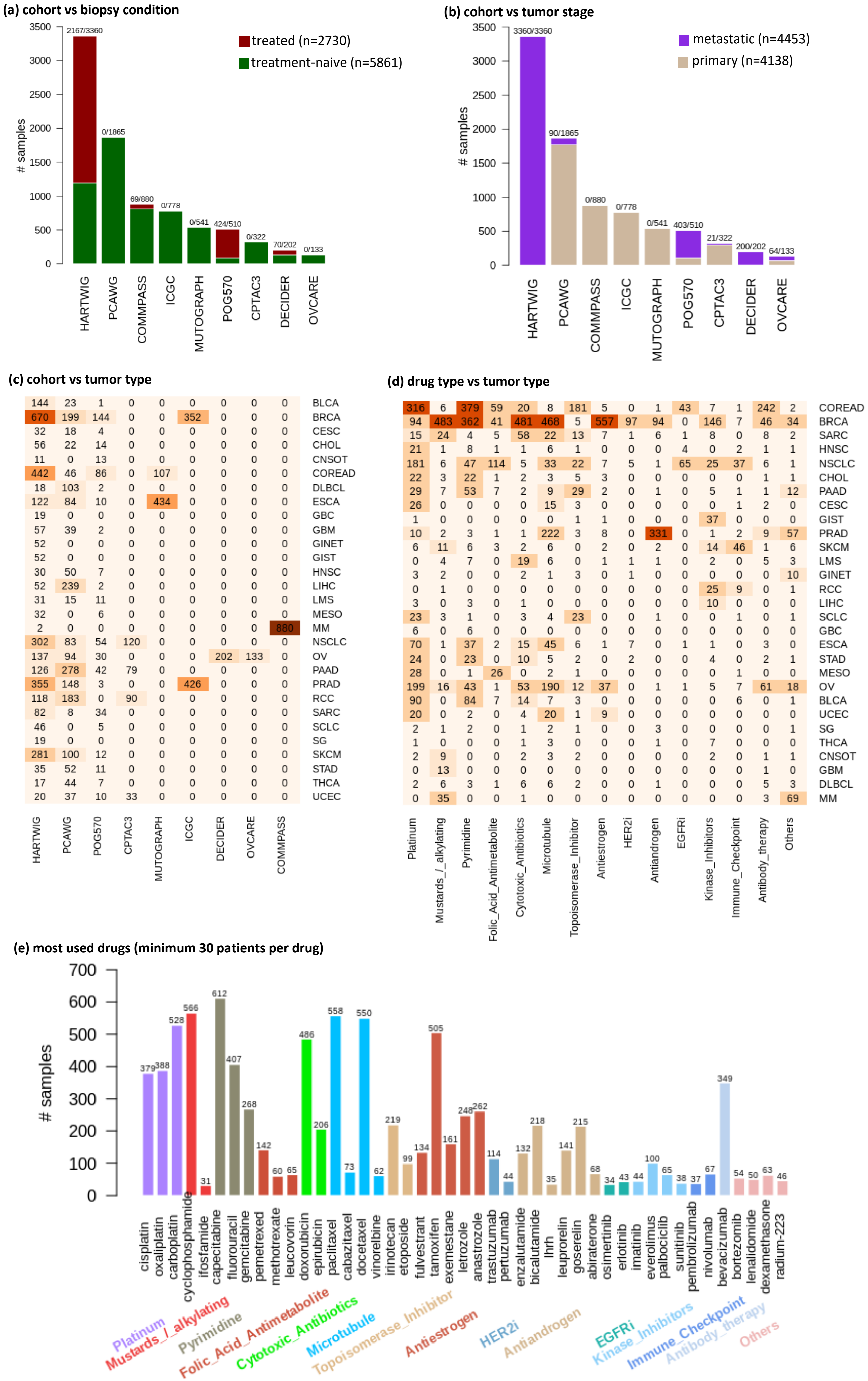
(a) cohort vs biopsy condition
treated (n=2730)
treatment-naive (n=5861)
### samples
3500
3000
2500
2000
1500
1000
500
0
2167/3360
0/1865
69/880
0/778
0/541
424/510
0/322
70/202
0/133
HARTWIG
PCAWG
COMMPASS
ICGC
MUTOGRAPH
POG570
CPTAC3
DECIDER
OVCARE
(b) cohort vs tumor stage
metastatic (n=4453)
primary (n=4138)
### samples
3360/3360
90/1865
0/880
0/778
0/541
403/510
21/322
200/202
64/133
HARTWIG
PCAWG
COMMPASS
ICGC
MUTOGRAPH
POG570
CPTAC3
DECIDER
OVCARE
(c) cohort vs tumor type
144 23 1 0 0 0 0 0 0 BLCA
670 199 144 0 0 352 0 0 0 BRCA
32 18 4 0 0 0 0 0 0 CESC
56 22 14 0 0 0 0 0 0 CHOL
11 0 13 0 0 0 0 0 0 CNSOT
442 46 86 0 107 0 0 0 0 COREAD
18 103 2 0 0 0 0 0 0 DLBCL
122 84 10 0 434 0 0 0 0 ESCA
19 0 0 0 0 0 0 0 0 GBC
57 39 2 0 0 0 0 0 0 GBM
52 0 0 0 0 0 0 0 0 GINET
52 0 0 0 0 0 0 0 0 GIST
30 50 7 0 0 0 0 0 0 HNSC
52 239 2 0 0 0 0 0 0 LIHC
31 15 11 0 0 0 0 0 0 LMS
32 0 6 0 0 0 0 0 0 MESO
2 0 0 0 0 0 0 0 880 MM
302 83 54 120 0 0 0 0 0 NSCLC
137 94 30 0 0 0 202 133 0 OV
126 278 42 79 0 0 0 0 0 PAAD
355 148 3 0 0 426 0 0 0 PRAD
118 183 0 90 0 0 0 0 0 RCC
82 8 34 0 0 0 0 0 0 SARC
46 0 5 0 0 0 0 0 0 SCLC
19 0 0 0 0 0 0 0 0 SG
281 100 12 0 0 0 0 0 0 SKCM
35 52 11 0 0 0 0 0 0 STAD
17 44 7 0 0 0 0 0 0 THCA
20 37 10 33 0 0 0 0 0 UCEC
HARTWIG
PCAWG
POG570
CPTAC3
MUTOGRAPH
ICGC
DECIDER
OVCARE
COMPASS
(d) drug type vs tumor type
316 6 379 59 20 8 181 5 0 1 43 7 1 242 2 COREAD
94 483 362 41 481 468 5 557 97 94 0 146 7 46 34 BRCA
15 24 4 5 58 22 13 7 1 6 1 8 0 8 2 SARC
21 1 8 1 1 6 1 0 0 0 4 0 2 1 1 HNSC
181 6 47 114 5 33 22 7 5 1 65 25 37 6 1 NSCLC
22 3 22 1 2 3 5 3 0 0 0 0 0 1 1 CHOL
29 7 53 7 2 9 29 2 0 1 0 5 1 1 12 PAAD
26 0 0 0 0 15 3 0 0 0 0 0 1 2 0 CESC
1 0 0 0 0 0 1 1 0 0 0 37 0 0 0 GIST
10 2 3 1 1 222 3 8 0 331 0 1 2 9 57 PRAD
6 11 6 3 2 6 0 2 0 2 0 14 46 1 6 SKCM
0 4 7 0 19 6 0 1 1 1 0 2 0 5 3 LMS
3 2 0 0 2 1 3 0 1 0 0 0 0 0 10 GINET
0 1 0 0 0 0 0 0 0 0 0 25 9 0 1 RCC
3 0 3 0 1 0 0 0 0 0 0 10 0 0 0 LIHC
23 3 1 0 3 4 23 0 0 1 0 0 1 0 1 SCLC
6 0 6 0 0 0 0 0 0 0 0 0 0 0 0 GBC
70 1 37 2 15 45 6 1 7 0 1 1 0 3 0 ESCA
24 0 23 0 10 5 2 0 2 0 0 4 0 2 1 STAD
28 0 1 26 0 2 1 0 0 0 0 0 1 0 0 MESO
199 16 43 1 53 190 12 37 0 1 1 5 7 61 18 OV
90 0 84 7 14 7 3 0 0 0 0 0 6 0 1 BLCA
20 0 2 0 4 20 0 9 0 0 0 0 0 0 0 UCEC
2 1 2 0 1 2 1 0 0 3 0 0 0 0 1 SG
1 0 0 0 1 3 0 0 0 1 0 7 0 0 0 THCA
2 9 0 0 2 3 2 0 0 0 0 2 0 1 1 CNSOT
0 13 0 0 0 0 0 0 0 0 0 0 0 1 0 GBM
2 6 3 1 6 6 2 0 0 1 0 0 0 5 3 DLBCL
0 35 0 0 1 0 0 0 0 0 0 0 0 3 69 MM
Platinum
Mustards_/_alkylating
Pyrimidine
Folic_Acid_Antimetabolite
Cytotoxic_Antibiotics
Microtubule
Topoisomerase_Inhibitor
Antiestrogen
HER2i
Antiandrogen
EGFRi
Kinase_Inhibitors
Immune_Checkpoint
Antibody_therapy
Others
(e) most used drugs (minimum 30 patients per drug)
### samples
700
600
500
400
300
200
100
0
cisplatin 379
oxaliplatin 388
carboplatin 528
cyclophosphamide 566
ifosfamide 31
capecitabine 612
fluorouracil 407
gemcitabine 268
pemetrexed 142
methotrexate 60
leucovorin 65
doxorubicin 486
epirubicin 206
paclitaxel 558
cabazitaxel 73
docetaxel 550
vinorelbine 62
irinotecan 219
etoposide 99
fulvestrant 134
tamoxifen 505
exemestane 161
letrozole 248
anastrozole 262
trastuzumab 114
pertuzumab 44
enzalutamide 132
bicalutamide 218
lhrh 35
leuprorelin 141
goserelin 215
abiraterone 68
osimertinib 34
erlotinib 43
imatinib 44
everolimus 100
palbociclib 65
sunitinib 38
pembrolizumab 37
nivolumab 67
bevacizumab 349
bortezomib 54
lenalidomide 50
dexamethasone 63
radium-223 46
Platinum
Mustards_/_alkylating
Pyrimidine
Folic_Acid_Antimetabolite
Cytotoxic_Antibiotics
Microtubule
Topoisomerase_Inhibitor
Antiestrogen
HER2i
Antiandrogen
EGFRi
Kinase_Inhibitors
Immune_Checkpoint
Antibody_therapy
Others

**Supplementary Figure 1. Overview of the dataset with additional information**. **a)** Number of treatment-naive and pre-treated samples from each study. **b)** bar plot showing the number of primary and metastatic samples from each study. **c)** Number of samples from each tumor type for each study. The x-axis represents the study and the y-axis represents the tumor type. **d)** Number of samples pre-treated with different drug types from each cancer type. The x-axis represents the drug type based on its mechanism of action and the y-axis represents the tumor type. **e)** The most frequent pre-treatment drugs within each mechanism-of-action category, showing drugs administered to >=30 patients.

**(a) differences in target and background mutations between metastatic patients pretreated with and without platinum-based drugs**

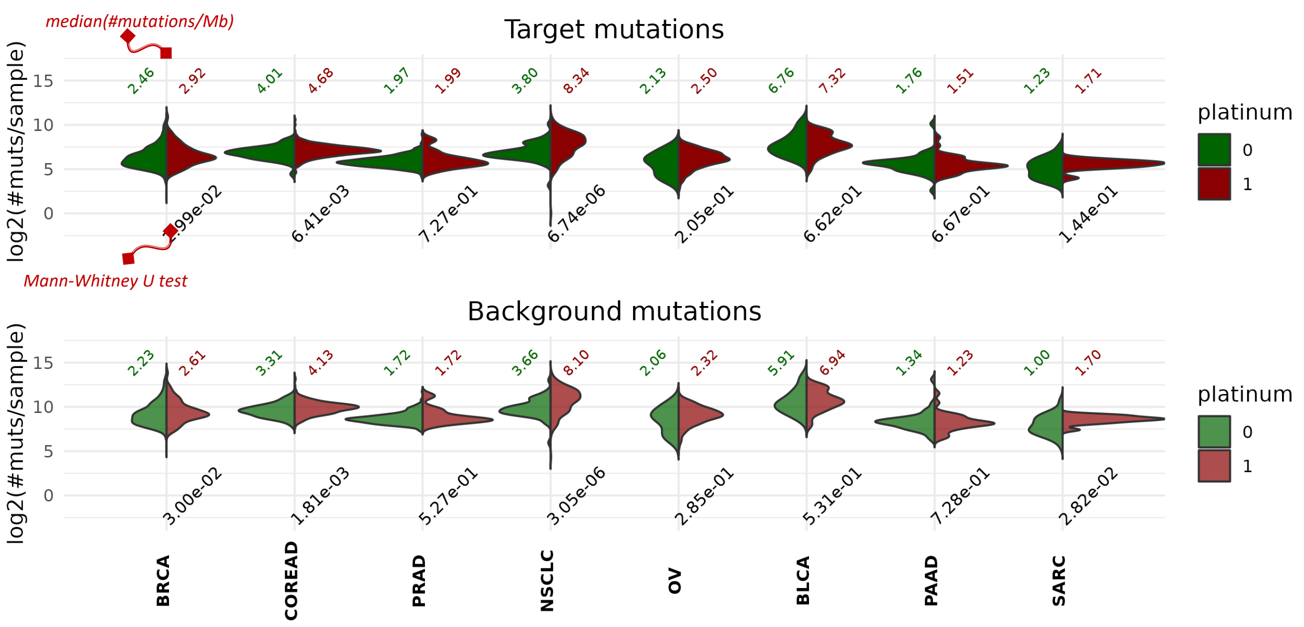


**(b) differences in target and background mutations between metastatic patients pretreated with and without pyrimidine drugs**

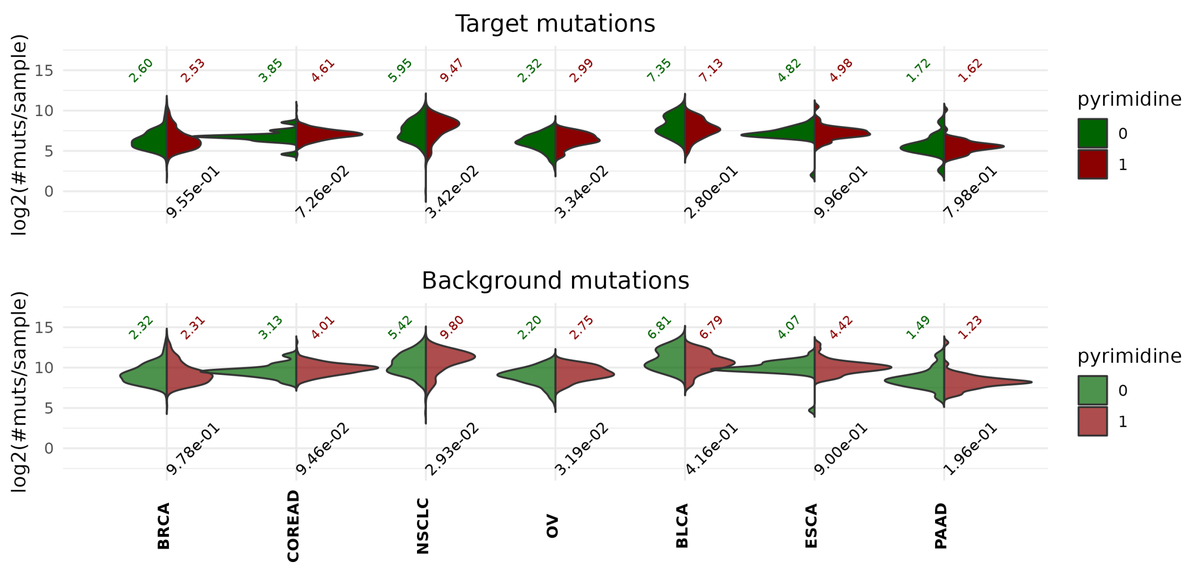


**(c) differences in target and background mutations between metastatic patients pretreated with and without microtubule drugs**

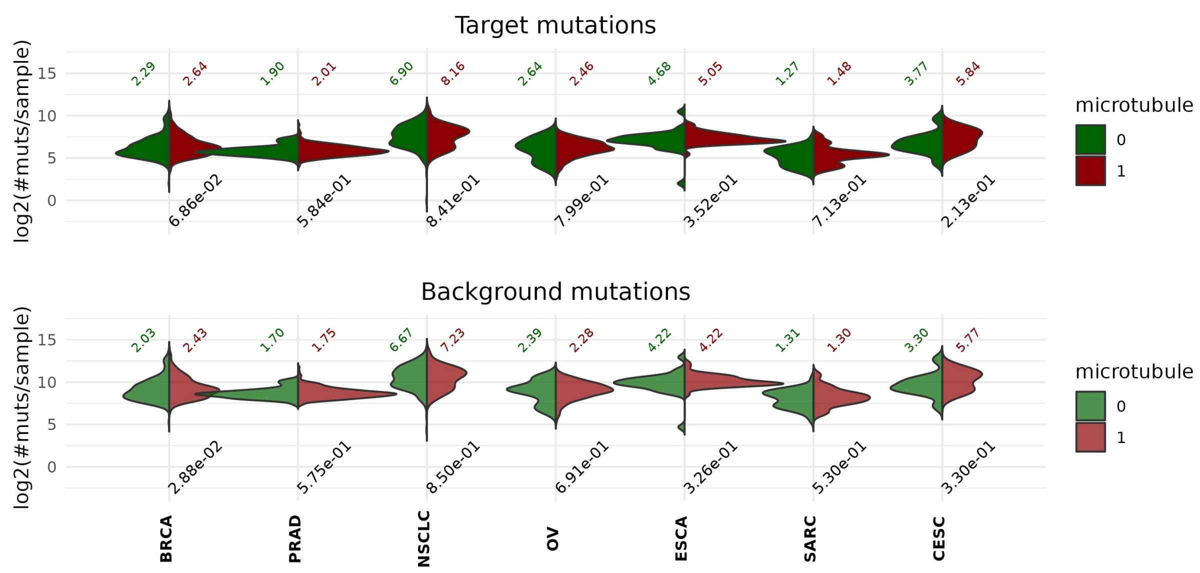

**Supplementary Figure 2. Changes in tumor mutation burden associated with specific drug types.** Distributions of the tumor mutation burden (here defined as number of point mutations per sample) between the tumors in metastatic patients pre-treated with a specific drug type (in red): **a)** platinum-based alkylating drugs, **b)** pyrimidine analog drugs, and **c)** microtubule drugs), contrasted against the metastatic patients pre-treated with other drugs (in green) in different cancer types. The top and bottom panels represent the changes in the exonic (DiffInvex target) regions and intronic/UTR/gene flanking (DiffInvex background) regions, respectively. The numbers on bottom of each panel are the p-values of the Mann-Whitney U test (two-tailed) that compare the tumor mutation burden per sample between the treatment-naive and pre-treated patients.

**(a) target mutation rate**

| Platinum | Mustards_/_alkylating | Pyrimidine | Folic_Acid_Antimetabolite | Cytotoxic_Antibiotics | Microtubule | Topoisomerase_Inhibitor | Antiestrogen | HER2i | Antiandrogen | EGFRi | Kinase_Inhibitors | Immune_Checkpoint | Antibody_therapy | Others | |
|---|---|---|---|---|---|---|---|---|---|---|---|---|---|---|---|
| 1.157 | 0.984 | 1.174 | 1.467 | 1.002 | 0.872 | 1.153 | 0.857 | 1.152 | 0.698 | 1.188 | 1.33 | 1.747 | 0.937 | 0.851 | Platinum |
| 0.984 | 0.893 | 0.947 | 1.239 | 0.833 | 0.853 | 0.654 | 0.862 | 0.993 | 0.651 | NaN | 0.942 | 3.782 | 1.14 | 0.666 | Mustards_/_alkylating |
| 1.174 | 0.947 | 1.064 | 1.181 | 1.018 | 0.906 | 0.975 | 0.889 | 1.527 | 0.738 | 1.152 | 0.96 | 1.486 | 0.935 | 1.124 | Pyrimidine |
| 1.467 | 1.239 | 1.181 | 1.424 | 1.059 | 1.378 | 1.28 | 1.101 | 3.54 | 0.676 | 1.266 | 1.517 | 2.356 | 0.972 | 1.521 | Folic_Acid_Antimetabolite |
| 1.002 | 0.833 | 1.018 | 1.059 | 0.859 | 0.859 | 0.951 | 0.862 | 1.153 | 0.737 | 0.998 | 1.002 | 1.034 | 0.742 | 0.958 | Cytotoxic_Antibiotics |
| 0.872 | 0.853 | 0.906 | 1.378 | 0.859 | 0.78 | 0.878 | 0.86 | 1.059 | 0.596 | 0.757 | 1.074 | 1.531 | 0.767 | 0.716 | Microtubule |
| 1.153 | 0.654 | 0.975 | 1.28 | 0.951 | 0.878 | 1.076 | 0.87 | 3.626 | 1.123 | 1.213 | 1.83 | 1.447 | 0.984 | 0.98 | Topoisomerase_Inhibitor |
| 0.857 | 0.862 | 0.889 | 1.101 | 0.862 | 0.86 | 0.87 | 0.826 | 1 | 0.665 | NaN | 0.96 | 0.786 | 0.827 | 1.255 | Antiestrogen |
| 1.152 | 0.993 | 1.527 | 3.54 | 1.153 | 1.059 | 3.626 | 1 | 1.049 | 1.284 | 0.931 | 3.082 | 0.076 | 1.147 | 2.778 | HER2i |
| 0.698 | 0.651 | 0.738 | 0.676 | 0.737 | 0.596 | 1.123 | 0.665 | 1.284 | 0.568 | NaN | 0.893 | 1.217 | 0.448 | 0.542 | Antiandrogen |
| 1.188 | NaN | 1.152 | 1.266 | 0.998 | 0.757 | 1.213 | NaN | 0.931 | NaN | 1.027 | 1.41 | 0.922 | 1.082 | NaN | EGFRi |
| 1.33 | 0.942 | 0.96 | 1.517 | 1.002 | 1.074 | 1.83 | 0.96 | 3.082 | 0.893 | 1.41 | 0.887 | 2.615 | 1.057 | 1.366 | Kinase_Inhibitors |
| 1.747 | 3.782 | 1.486 | 2.356 | 1.034 | 1.531 | 1.447 | 0.786 | 0.076 | 1.217 | 0.922 | 2.615 | 2.266 | 3.73 | 1.158 | Immune_Checkpoint |
| 0.937 | 1.14 | 0.935 | 0.972 | 0.742 | 0.767 | 0.984 | 0.827 | 1.147 | 0.448 | 1.082 | 1.057 | 3.73 | 0.913 | 0.502 | Antibody_therapy |
| 0.851 | 0.666 | 1.124 | 1.521 | 0.958 | 0.716 | 0.98 | 1.255 | 2.778 | 0.542 | NaN | 1.366 | 1.158 | 0.502 | 0.645 | Others |

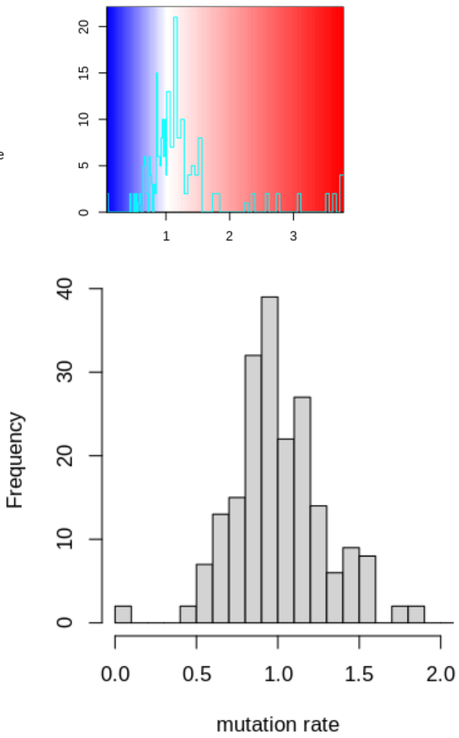


**(b) target/background mutation rate**

| Platinum | Mustards_/_alkylating | Pyrimidine | Folic_Acid_Antimetabolite | Cytotoxic_Antibiotics | Microtubule | Topoisomerase_Inhibitor | Antiestrogen | HER2i | Antiandrogen | EGFRi | Kinase_Inhibitors | Immune_Checkpoint | Antibody_therapy | Others | |
|---|---|---|---|---|---|---|---|---|---|---|---|---|---|---|---|
| 1.014 | 1.004 | 1.029 | 0.993 | 1.019 | 1.015 | 1.046 | 1.012 | 1.004 | 0.991 | 1.023 | 0.974 | 0.97 | 1.054 | 1.011 | Platinum |
| 1.004 | 0.988 | 0.987 | 0.989 | 0.996 | 0.995 | 1.028 | 0.996 | 0.979 | 1.01 | NaN | 0.989 | 0.934 | 0.962 | 1.001 | Mustards_/_alkylating |
| 1.029 | 0.987 | 1.024 | 1.02 | 1.001 | 0.987 | 1.09 | 0.998 | 0.953 | 1.011 | 1.034 | 1.004 | 0.959 | 1.069 | 0.997 | Pyrimidine |
| 0.993 | 0.989 | 1.02 | 0.991 | 0.997 | 0.985 | 1.003 | 0.999 | 0.894 | 1.04 | 0.998 | 0.981 | 0.969 | 1.047 | 1.026 | Folic_Acid_Antimetabolite |
| 1.019 | 0.996 | 1.001 | 0.997 | 1.005 | 0.994 | 1.106 | 0.996 | 0.967 | 0.999 | 1.088 | 0.987 | 0.968 | 0.999 | 1.014 | Cytotoxic_Antibiotics |
| 1.015 | 0.995 | 0.987 | 0.985 | 0.994 | 1.009 | 1 | 0.998 | 0.983 | 1.009 | 1.1 | 0.989 | 0.98 | 0.994 | 1.028 | Microtubule |
| 1.046 | 1.028 | 1.09 | 1.003 | 1.106 | 1 | 1.053 | 1.02 | 0.835 | 0.972 | 1.029 | 0.954 | 1.069 | 1.083 | 1.005 | Topoisomerase_Inhibit |
| 1.012 | 0.996 | 0.998 | 0.999 | 0.996 | 0.998 | 1.02 | 1.003 | 0.985 | 1.003 | NaN | 1.003 | 1.011 | 1.006 | 1.018 | Antiestrogen |
| 1.004 | 0.979 | 0.953 | 0.894 | 0.967 | 0.983 | 0.835 | 0.985 | 0.985 | 0.943 | 1.127 | 0.961 | 1.011 | 1.043 | 0.963 | HER2i |
| 0.991 | 1.01 | 1.011 | 1.04 | 0.999 | 1.009 | 0.972 | 1.003 | 0.943 | 1.02 | NaN | 0.987 | 1.028 | 1.027 | 1.015 | Antiandrogen |
| 1.023 | NaN | 1.034 | 0.998 | 1.088 | 1.1 | 1.029 | NaN | 1.127 | NaN | 1.024 | 0.966 | 1.119 | 1.019 | NaN | EGFRi |
| 0.974 | 0.989 | 1.004 | 0.981 | 0.987 | 0.989 | 0.954 | 1.003 | 0.961 | 0.987 | 0.966 | 0.993 | 0.961 | 0.969 | 0.988 | Kinase_Inhibitors |
| 0.97 | 0.934 | 0.959 | 0.969 | 0.968 | 0.98 | 1.069 | 1.011 | 1.011 | 1.028 | 1.119 | 0.961 | 0.959 | 0.908 | 0.899 | Immune_Checkpoint |
| 1.054 | 0.962 | 1.069 | 1.047 | 0.999 | 0.994 | 1.083 | 1.006 | 1.043 | 1.027 | 1.019 | 0.969 | 0.908 | 1.05 | 1.02 | Antibody_therapy |
| 1.011 | 1.001 | 0.997 | 1.026 | 1.014 | 1.028 | 1.005 | 1.018 | 0.963 | 1.015 | NaN | 0.988 | 0.899 | 1.02 | 0.99 | Others |

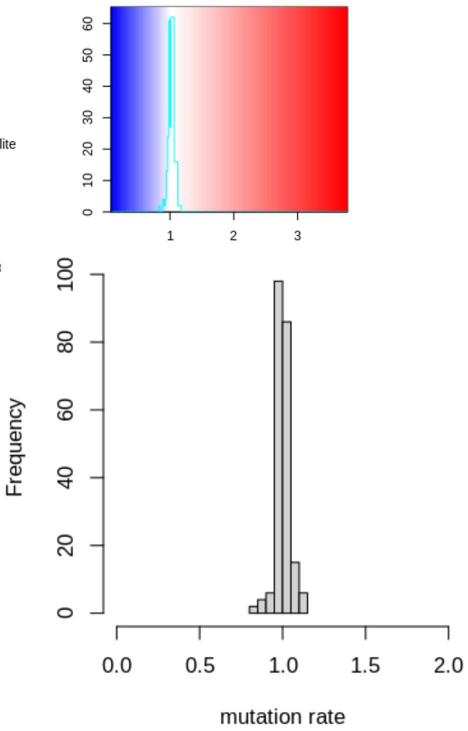

**Supplementary Figure 3. Changes in tumor mutation burden associated with different combinations of drug types. a)** Comparison between the DiffInvex target region (coding exonic) mutation rate per sample, divided by the median mutation rate per sample, in patients pre-treated with different drug combinations. Each element in the heatmap shows the target region mutation rate per sample of patients pre-treated with the combination of two drug types on the x-axis and y-axis, with or without other drug types. Elements on the diagonal represent the target region mutation rate of patients pre-treated with the drug type on the x-axis, with or without other drug types. The histogram shows the distribution of the normalized target region mutation rates per sample, across samples pre-treated with different drug combinations. **b)** Comparison between the ratio of DiffInvex target region to DiffInvex background region (introns, UTRs, gene flanks) mutations per sample, divided by their median ratio, in patients pre-treated with different drug combinations. Elements in the heatmap as in panel **a**. The histogram shows the distribution of the normalized ratios of target mutations to background mutations across samples pre-treated with different drug combinations.

(a) Observed versus estimated exonic mutations for different tumor types

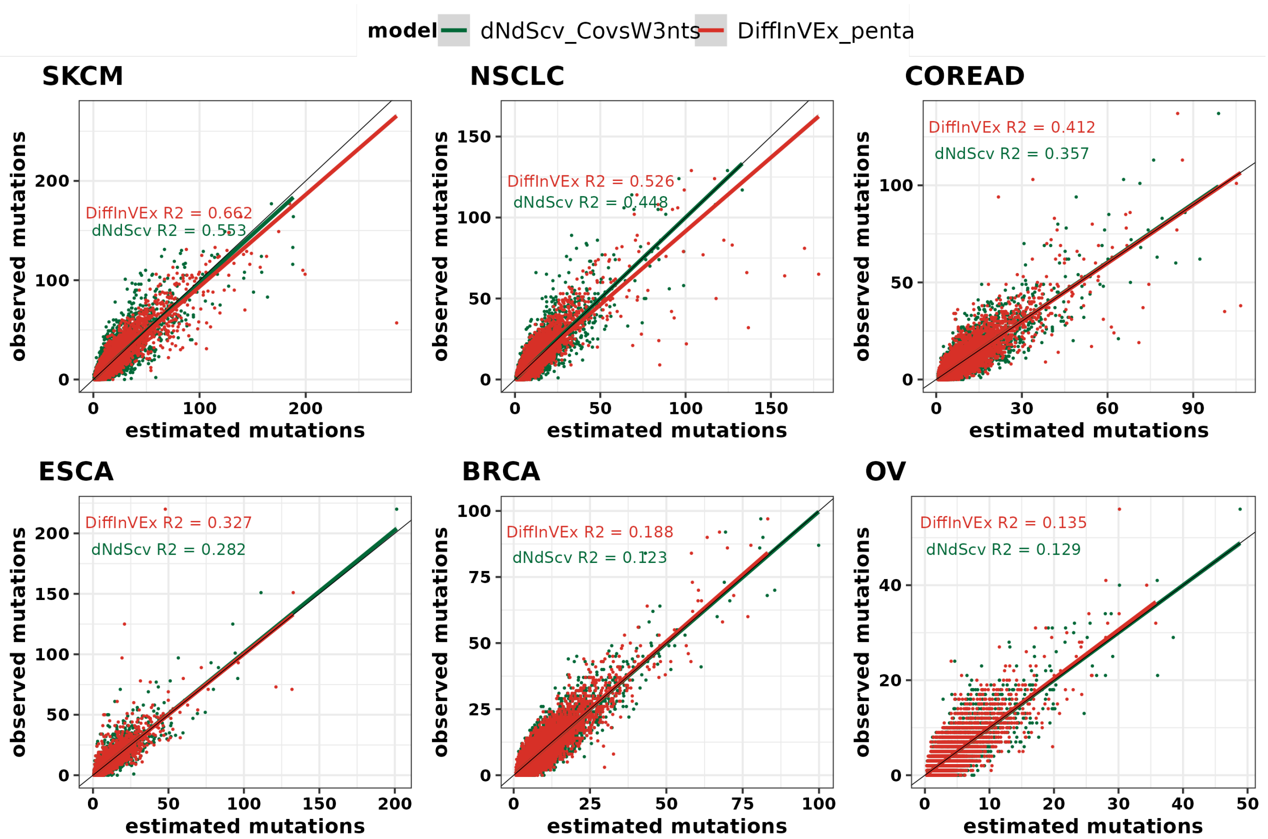


(b) Performance evaluation of DiffInvex and dNdScv methods to identify cancer drivers

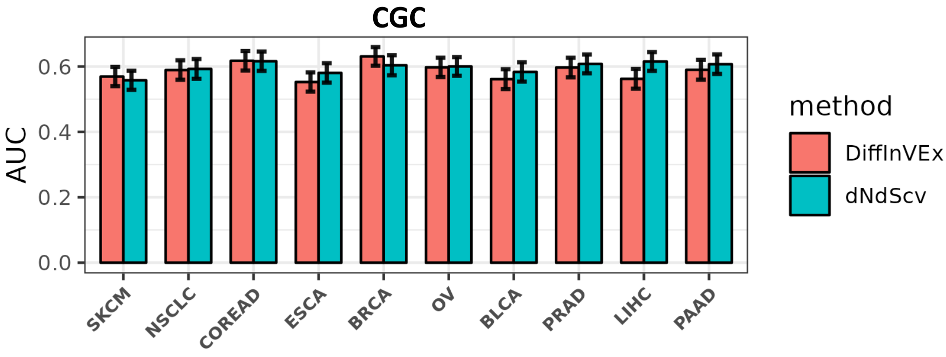

**Supplementary Figure 4. Benchmarking the DiffInvex framework by analyses of somatic selection in a general, not conditional, setting. a)** comparison between the DiffInvex intronic rate baseline, and the dNdScv covariate-based baselines in explaining the variation in coding mutation rate for non-cancerous genes in different tumor types. Plots show agreement between the estimated and observed mutation counts using the DiffInvex with pentanucleotide matching (in red) and the dNdScv covariates additionally supported with trinucleotide composition (in green). **b)** comparison of the accuracy (as AUC, Area under the ROC Curve) between DiffInvex and dNdScv in identifying known cancer genes in Cancer Gene Census (CGC) from a background set of likely passenger genes.

(a) Comparsion between the mutation baseline between DiffInvex versions

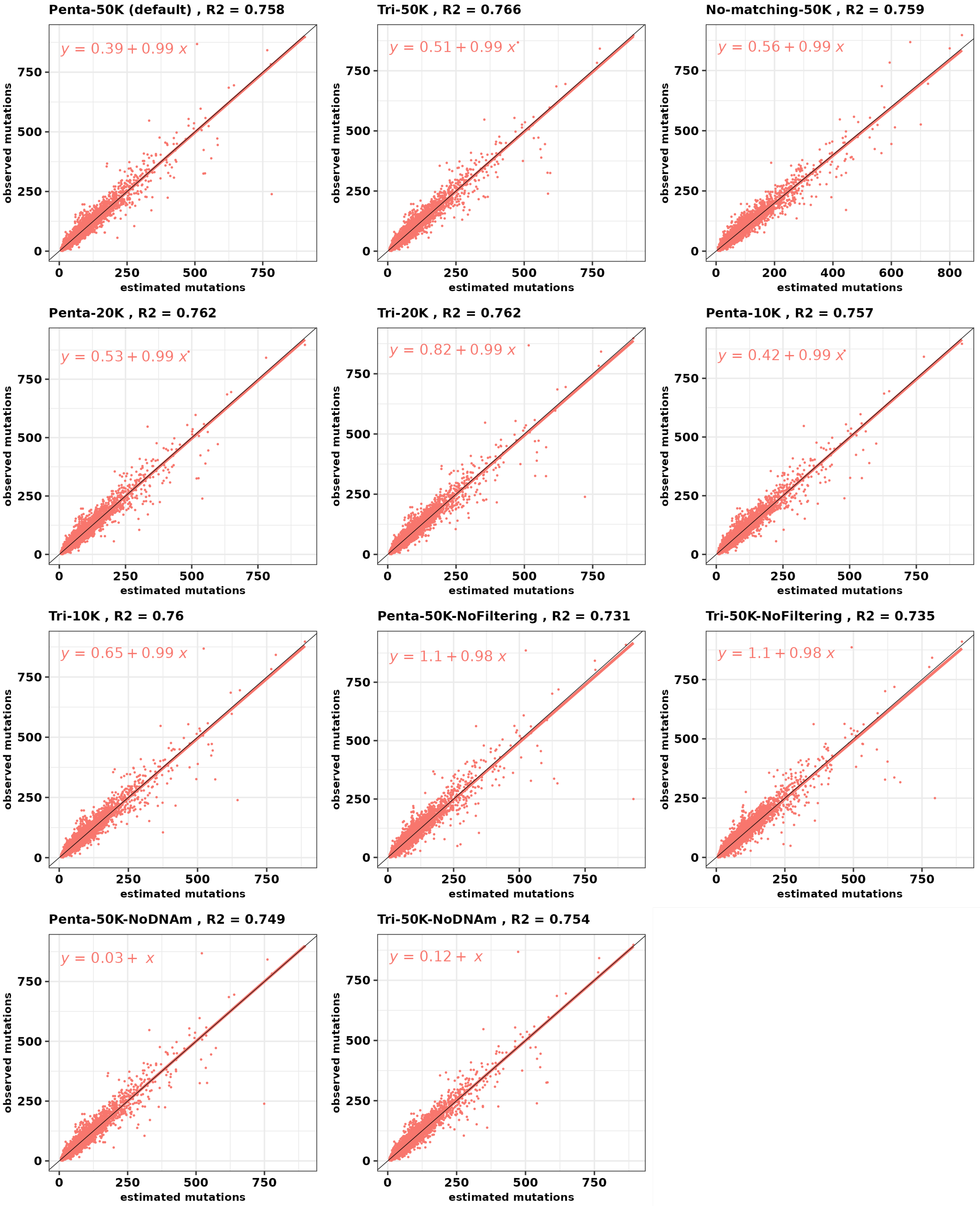

**Supplementary Figure 5. Evaluation of DiffInvex method variations on explaining the variations in exonic coding mutation rate.** Each panel shows the scatter plot between the observed mutation counts and the estimated mutations in passenger genes in a pan-cancer analysis, using a different setup of DiffInvex as stated on the top of each panel. **Penta-50K**: the default DiffInvex setup using 50,000 nucleotides as background region minimum width (before matching), implementing all filters for problematic genomic regions (see Methods for details) and matching the pentanucleotide and DNA-methylated sites composition between target (exonic) regions and background (intronic, flanks and UTRs) regions. **Tri-50K**: same as penta-50K setup, but matching based on the trinucleotide compositions only. **No-matching-50K**: uses 50,000 nucleotides as background width and implements all filters, without matching tri/pentanucleotides between target and background regions. **Penta-20K and Penta-10K**: same as Penta-50K setup, but using 20,000 and 10,000 nucleotides as background width before matching, respectively. **Tri-20K and Tri-10K**: same as Tri-50K setup, but using 20,000 and 10,000 nucleotides as background width before matching, respectively. **Penta-50K-NoFiltering**: same as Penta-50K setup but without implementing any filter. **Tri-50K-NoFiltering**: same as Tri-50K setup but without implementing any filter. **Penta-50K-NoDNAm**: same as Penta-50K setup but without matching for the composition of DNA-methylated sites. **Tri-50K-NoDNAm**: same as Tri-50K setup but without matching for the composition of DNA-methylated sites. The R2 values are stated next to the model name.

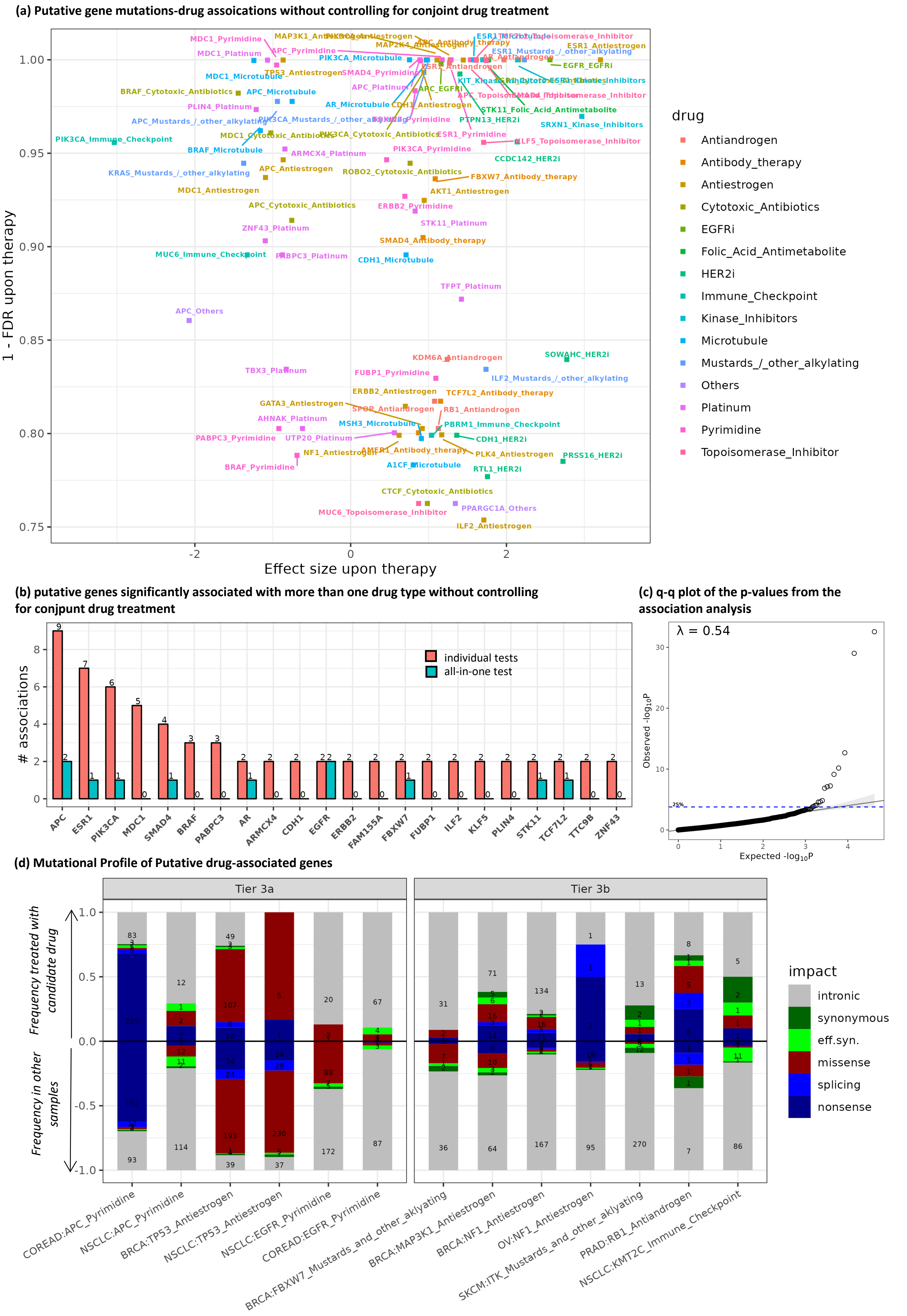
(a) Putative gene mutations-drug assoications without controlling for conjoint drug treatment
1 - FDR upon therapy
Effect size upon therapy
drug
Antiandrogen
Antibody_therapy
Antiestrogen
Cytotoxic_Antibiotics
EGFRi
Folic_Acid_Antimetabolite
HER2i
Immune_Checkpoint
Kinase_Inhibitors
Microtubule
Mustards_/_other_alkylating
Others
Platinum
Pyrimidine
Topoisomerase_Inhibitor
PIK3CA_Immune_Checkpoint
APC_Others
TFPT_Platinum
SOWAHC_HER2i
KDM6A_Antiandrogen
ILF2_Mustards_/_other_alkylating
TCF7L2_Antibody_therapy
RB1_Antiandrogen
PBRM1_Immune_Checkpoint
CDH1_HER2i
PLK4_Antiestrogen
PRSS16_HER2i
RTL1_HER2i
CTCF_Cytotoxic_Antibiotics
PPARGC1A_Others
MUC6_Topoisomerase_Inhibitor
ILF2_Antiestrogen
(b) putative genes significantly associated with more than one drug type without controlling for conjpunt drug treatment
### associations
individual tests
all-in-one test
APC
ESR1
PIK3CA
MDC1
SMAD4
BRAF
PABPC3
AR
ARMCX4
CDH1
EGFR
ERBB2
FAM155A
FBXW7
FUBP1
ILF2
KLF5
PLIN4
STK11
TCF7L2
TTC9B
ZNF43
(c) q-q plot of the p-values from the association analysis
λ = 0.54
Observed -log10P
Expected -log10P
(d) Mutational Profile of Putative drug-associated genes
Tier 3a
Tier 3b
Frequency treated with candidate drug
Frequency in other samples
impact
intronic
synonymous
eff.syn.
missense
splicing
nonsense
COREAD:APC_Pyrimidine
NSCLC:APC_Pyrimidine
BRCA:TP53_Antiestrogen
NSCLC:TP53_Antiestrogen
NSCLC:EGFR_Pyrimidine
COREAD:EGFR_Pyrimidine
BRCA:FBXW7_Mustards_and_other_aklyating
BRCA:MAP3K1_Antiestrogen
BRCA:NF1_Antiestrogen
OV:NF1_Antiestrogen
SKCM:ITK_Mustards_and_other_aklyating
PRAD:RB1_Antiandrogen
NSCLC:KMT2C_Immune_Checkpoint

**Supplementary Figure 6: Chemotherapy-associated mutational drivers genes identified by DiffInvex without controlling for conjoint drug treatment. a)** putative drug sensitive and resistant associations identified in the pan-cancer analysis of 8,591 cancer genomes by comparing the mutation profiles of patients exposed to a specific group of drugs. The x-axis represents the effect size upon treatment, the natural log fold-enrichment of coding exonic mutation rates in tumor samples pre-treated with a certain drug type compared to treatment-naive samples, normalizing for the change in intronic mutation burdens. **b)** Comparing the number of associations with each gene without (in red) and with controlling (in blue) for conjoint drug treatments, suggesting that not controlling results in many spurious associations. **c)** Quantile-quantile plot of the p-values from the pan-cancer analysis by DiffInvex, with controlling for the conjoint drug treatment, suggesting that p-values are conservative. **d)** The mutational profile of putative drug-associated genes in candidate tumor types. The upper-half of the box plot (positive values) represents the fractions of different mutation types ( nonsense, splicing, high-impact missense, effectively synonymous (low-impact missense), synonymous, and intronic) in the candidate gene in tumor samples treated with the given drug type, while the lower-half of the box plot (negative values) represents the fractions of these mutation types in that gene in other tumor samples (treatment-naive samples and samples pre-treated with other drug types). The effectively synonymous ("eff.syn.") mutations are low-impact missense mutations that have AlphaMissense pathogenicity score < 0.39. Tier 3a are putative gene mutation-drug associations (FDR<=25% in the pan-cancer analysis) of a known cancer, but these associations indicate different responses (sensitivity or resistance) in different tumor types. Tier 3b has candidate tumor types of more tentative new gene mutation-drug associations (FDR <= 80% in the pan-cancer analysis) that could be supported by the functional impact test (Fig. 4a) and/or in independent panel-sequencing cohorts (Fig. 4b).

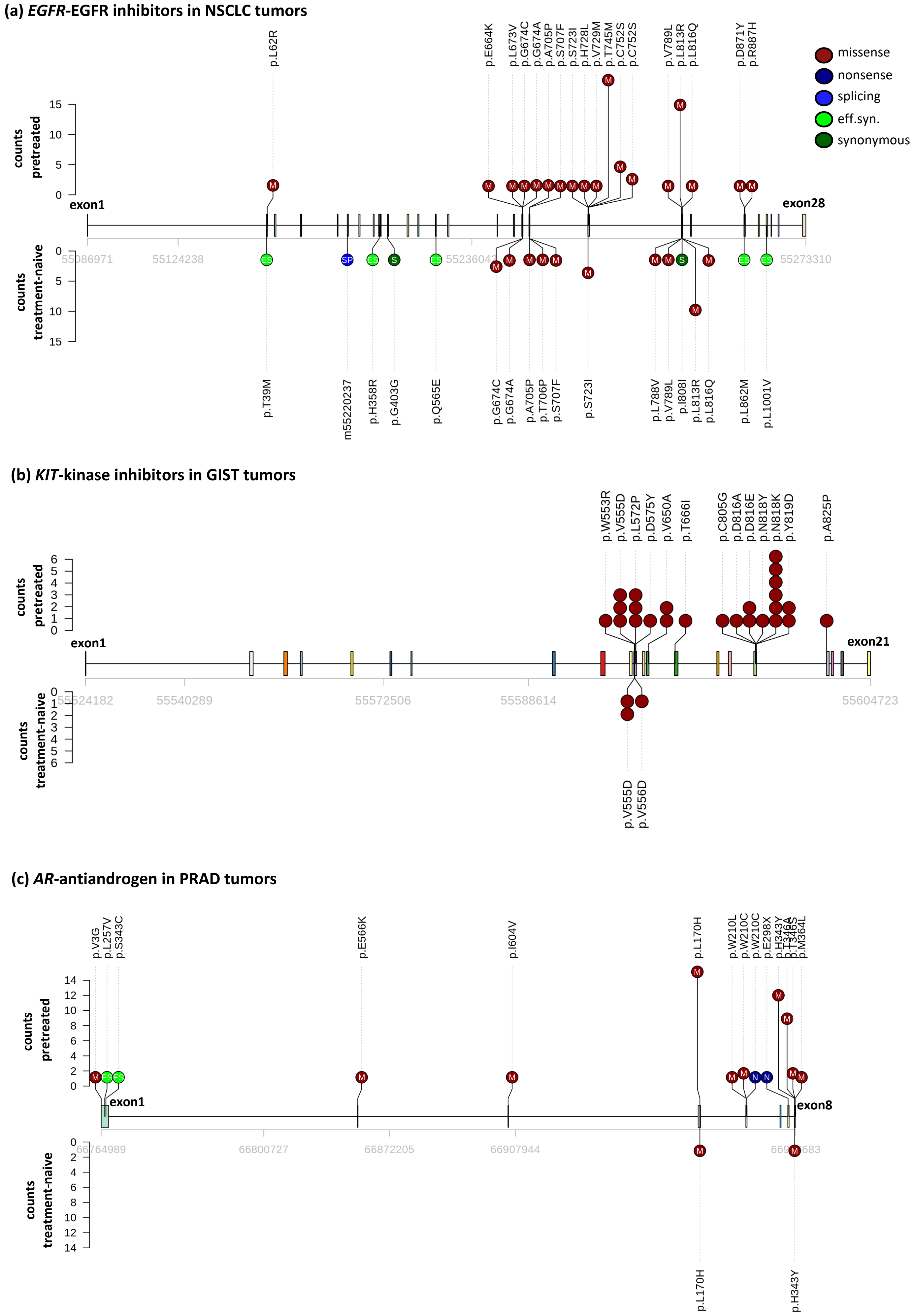
(a) EGFR-EGFR inhibitors in NSCLC tumors
missense
nonsense
splicing
eff.syn.
synonymous
counts pretreated
counts treatment-naive
exon1
exon28
55086971
55124238
55236042
55273310
p.L62R
p.E664K
p.L673V
p.G674C
p.G674A
p.A705P
p.S707F
p.S723I
p.H728L
p.V729M
p.T745M
p.C752S
p.C752S
p.V789L
p.L813R
p.L816Q
p.D871Y
p.R887H
p.T39M
m55220237
p.H358R
p.G403G
p.Q565E
p.G674C
p.G674A
p.A705P
p.T706P
p.S707F
p.S723I
p.L788V
p.V789L
p.I808I
p.L813R
p.L816Q
p.L862M
p.L1001V
(b) KIT-kinase inhibitors in GIST tumors
exon1
exon21
55524182
55540289
55572506
55588614
55604723
p.W553R
p.V555D
p.L572P
p.D575Y
p.V650A
p.T666I
p.C805G
p.D816A
p.D816E
p.N818Y
p.N818K
p.Y819D
p.A825P
p.V555D
p.V556D
(c) AR-antiandrogen in PRAD tumors
exon1
exon8
66764989
66800727
66872205
66907944
p.V3G
p.L257V
p.S343C
p.E566K
p.I604V
p.L170H
p.W210L
p.W210C
p.W210C
p.E298X
p.H343Y
p.M364L
p.L170H
p.H343Y

**Supplementary Figure 7. Distribution of exonic mutations across the gene body of Tier 1 genes.** Comparsion of the distribution of individual mutations in known drug resistance genes between the pre-treated and treatment-naive tumors for the putative associations: **a)** *EGFR*-EGFR inhibitors in NSCLC, **b)** *KIT*-kinase inhibitors in GIST and **c**) *AR*-Antiandrogen drugs in PRAD. In each panel, the upper-half of the lollipop plot shows the distribution of different mutation types (high-impact missense, nonsense, splicing, effectively synonymous (low-impact missense), and synonymous) in the candidate gene in tumor samples treated with the given drug type, while the lower-half of the lolliplot plot shows the distribution of different mutation types (high-impact missense, nonsense, splicing, effectively synonymous (low-impact missense), and synonymous) in the treatment-naive samples. The effectively synonymous ("eff.syn.") mutations are low-impact missense mutations that have AlphaMissense pathogenicity score < 0.39.

**(a) *PIK3CA*-antiestrogen drugs in BRCA tumors**

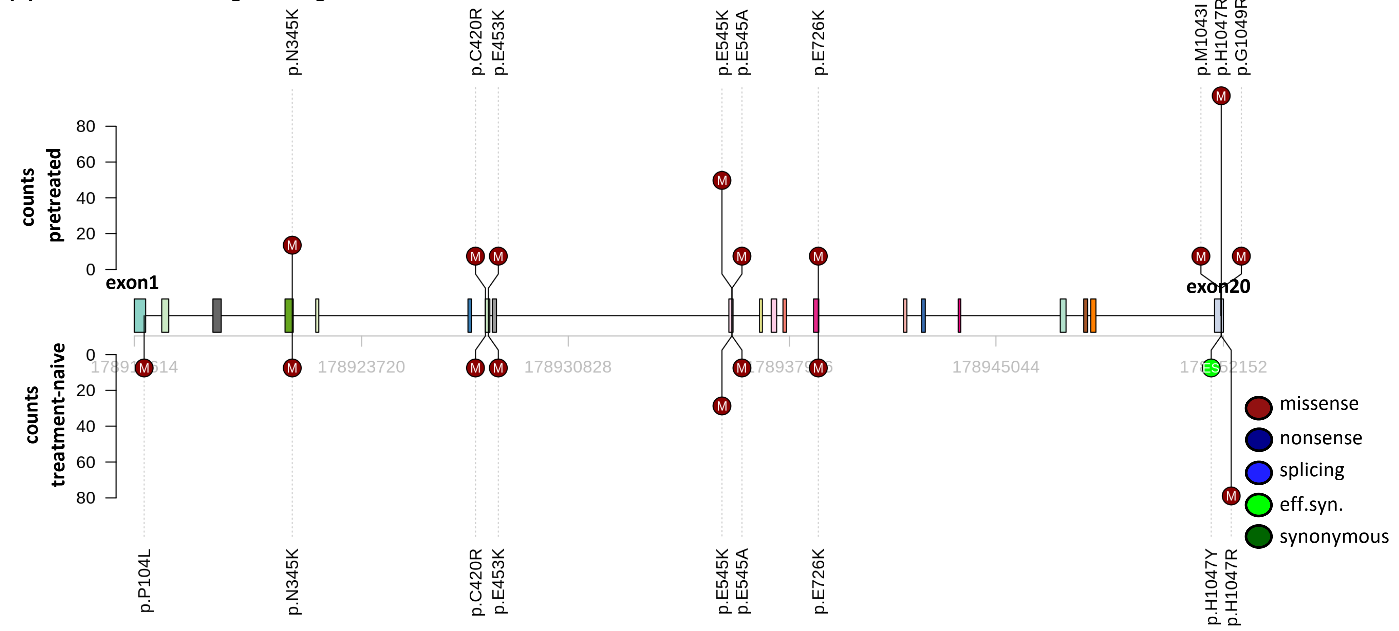


**(b) *APC*-antibody therapy in COREAD tumors**

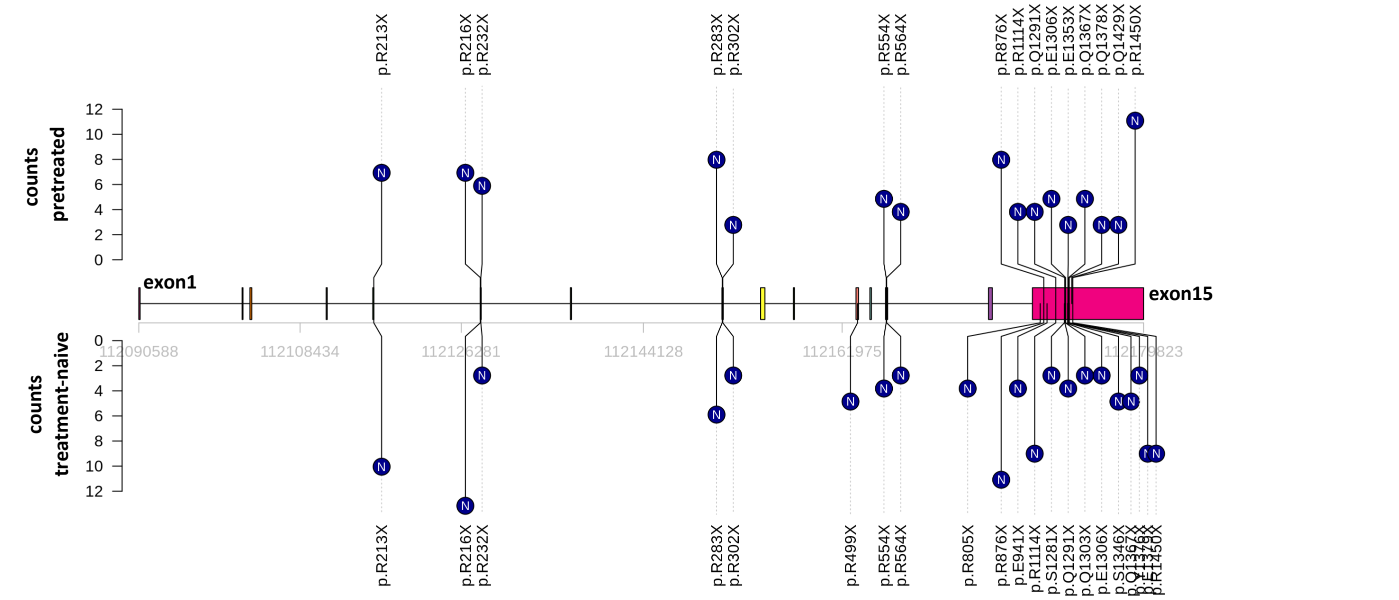


**(c) *STK11*-folic acid antimetabolite in NSCLC tumors**

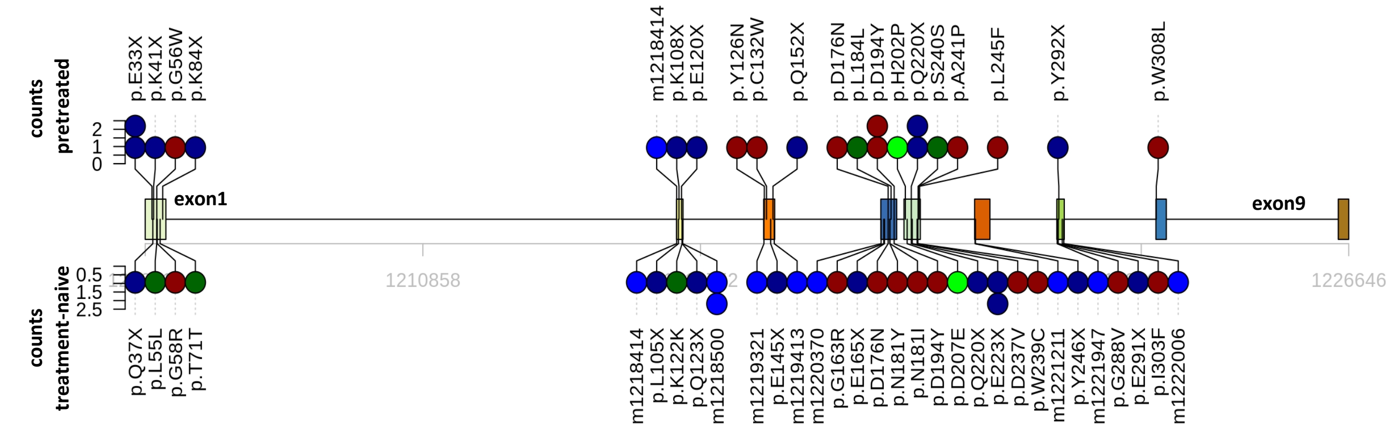

**Supplementary Figure 8. Distribution of exonic mutations across the gene body of Tier 2 genes.** Comparsion of the distribution of individual mutations in known drug resistance genes between the pre-treated and treatment-naive tumors for the putative associations: **a)** *PIK3CA*-antiestrogen drugs in BRCA, **b)** *APC*-antibody therapy in COREAD and **c**) *STK11*-folic acid antimetabolite drugs in NSCLC. In each panel, the upper-half of the lollipop plot shows the distribution of different mutation types (high-impact missense, nonsense, splicing, effectively synonymous (low-impact missense), and synonymous) in the candidate gene in tumor samples treated with the given drug type, while the lower-half of the lolliplot plot shows the distribution of different mutation types (high-impact missense, nonsense, splicing, effectively synonymous (low-impact missense), and synonymous) in the treatment-naive samples. The effectively synonymous ("eff.syn.") mutations are low-impact missense mutations that have AlphaMissense pathogenicity score < 0.39. In panels **a)** and **b)**, we showed only genomic positions with more than two mutations.

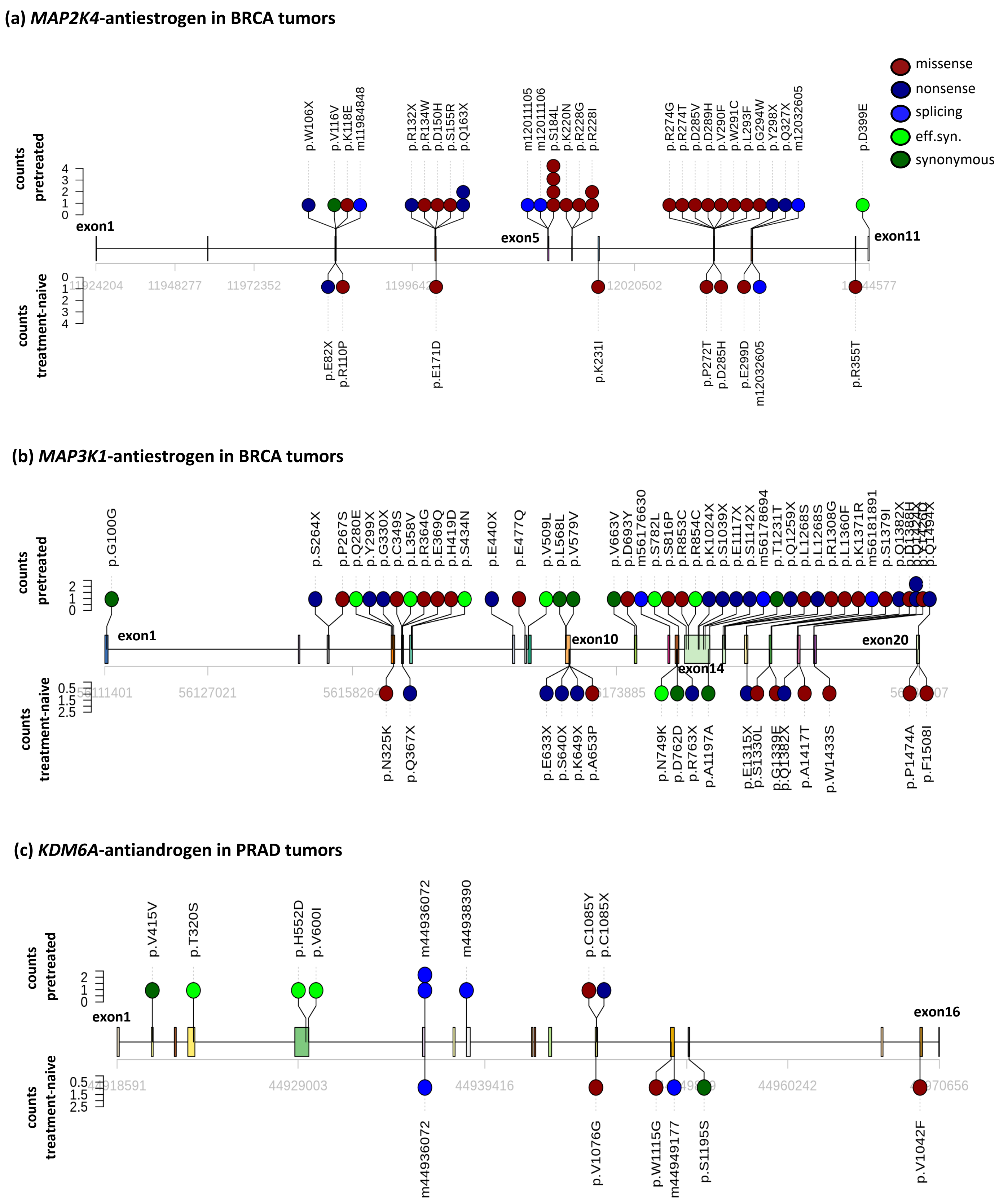

(a) MAP2K4-antiestrogen in BRCA tumors
missense
nonsense
splicing
eff.syn.
synonymous
counts pretreated
counts treatment-naive
exon1
exon5
exon11
p.W106X
p.V116V
p.K118E
m11984848
p.R132X
p.R134W
p.D150H
p.S155R
p.Q163X
m12011105
m12011106
p.S184L
p.K220N
p.R228G
p.R228I
p.R274G
p.R274T
p.D285V
p.D289H
p.V290F
p.W291C
p.L293F
p.G294W
p.Y298X
p.Q327X
m12032605
p.D399E
11924204
11948277
11972352
12020502
p.E82X
p.R110P
p.E171D
p.K231I
p.P272T
p.D285H
p.E299D
m12032605
p.R355T
(b) MAP3K1-antiestrogen in BRCA tumors
counts pretreated
counts treatment-naive
exon1
exon10
exon14
exon20
p.G100G
p.S264X
p.P267S
p.Q280E
p.Y299X
p.G330X
p.C349S
p.L358V
p.R364G
p.E369Q
p.H419D
p.S434N
p.E440X
p.E477Q
p.V509L
p.L568L
p.V579V
p.V663V
p.D693Y
m56176630
p.S782L
p.S816P
p.R853C
p.R854C
p.K1024X
p.S1039X
p.E1117X
p.S1142X
m56178694
p.T1231T
p.Q1259X
p.L1268S
p.L1268S
p.R1308G
p.L1360F
p.K1371R
m56181891
p.S1379I
p.Q1382X
56127021
p.N325K
p.Q367X
p.E633X
p.S640X
p.K649X
p.A653P
p.N749K
p.D762D
p.R763X
p.A1197A
p.E1315X
p.S1330L
p.A1417T
p.W1433S
p.P1474A
p.F1508I
(c) KDM6A-antiandrogen in PRAD tumors
counts pretreated
counts treatment-naive
exon1
exon16
p.V415V
p.T320S
p.H552D
p.V600I
m44936072
m44938390
p.C1085Y
p.C1085X
44929003
44939416
44960242
m44936072
p.V1076G
p.W1115G
m44949177
p.S1195S
p.V1042F

**Supplementary Figure 9. Distribution of exonic mutations across the gene body of Tier 2 and Tier 3 genes.** Comparsion of the distribution of individual mutations in known drug resistance genes between the pre-treated and treatment-naive tumors for the putative associations: **a)** *MAP2K4*-antiestrogen in BRCA, **b)** *MAP3K1*-antiestrogen in BRCA and **c**) *KDM6A*-antiandrogen drugs in PRAD. In each panel, the upper-half of the lollipop plot shows the distribution of different mutation types (high-impact missense, nonsense, splicing, effectively synonymous (low-impact missense), and synonymous) in the candidate gene in tumor samples treated with the given drug type, while the lower-half of the lolliplot plot shows the distribution of different mutation types (high-impact missense, nonsense, splicing, effectively synonymous (low-impact missense), and synonymous) in the treatment-naive samples. The effectively synonymous ("eff.syn.") mutations are low-impact missense mutations that have AlphaMissense pathogenicity score < 0.39.

**(a) Correlation between dEXdIn and dNdES ratios of cancer genes**

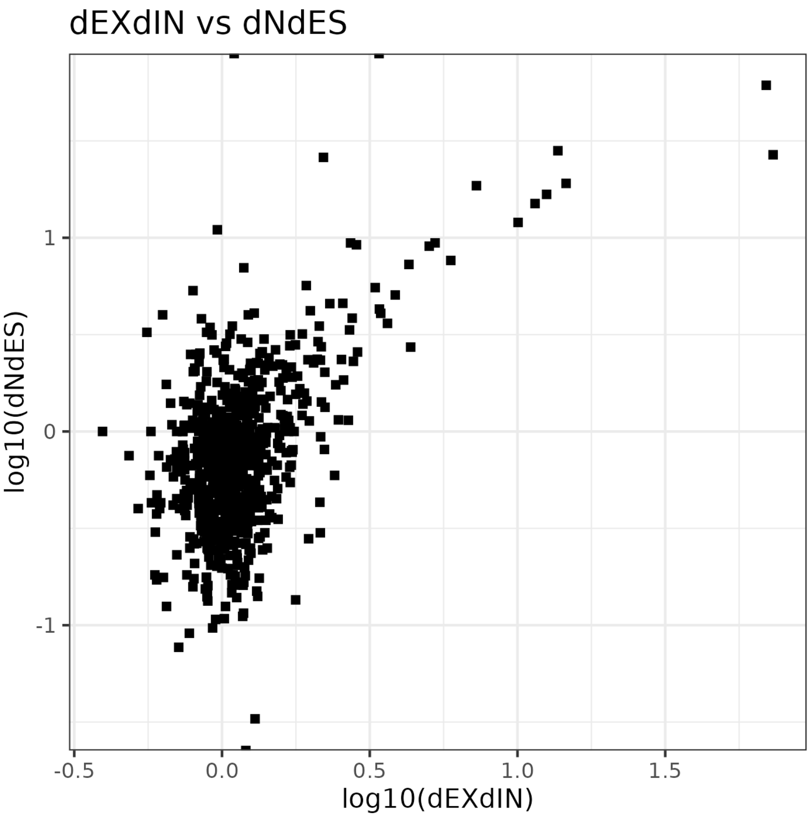


**(b) Distributions of dEXdIn and dNdES ratios of cancer genes**

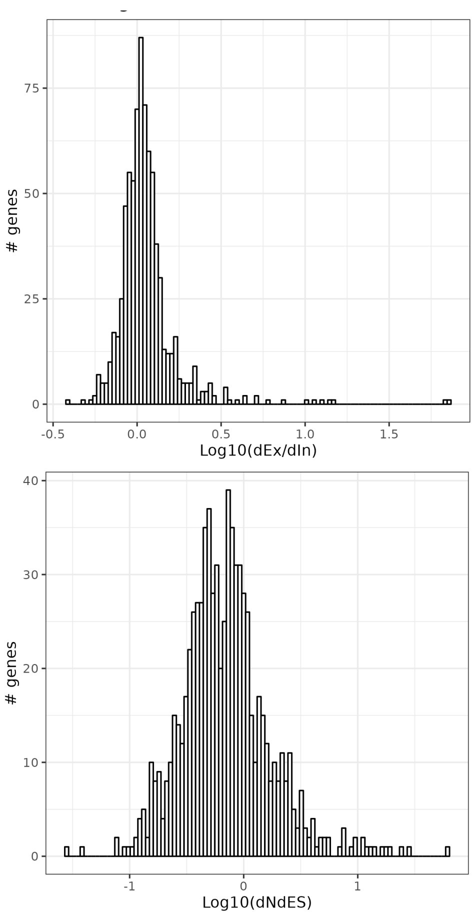


**(c) Slopes of linear regression between dEXdIn and dNdES ratios of cancer genes at different AlphaMissense pathogenic thersholds**

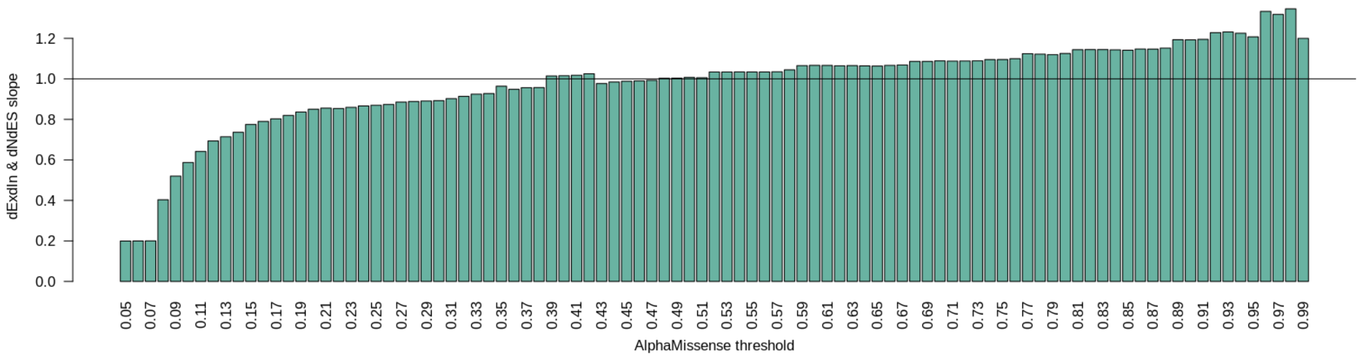

**Supplementary Figure 10. The dNdES test for conditional selection via differential functional impact. a)** Correlation between the effect sizes from the DiffInvex test (here, log10 of the ratio of exonic to intronic mutations, labeled dEXdIN) and the effect size from the dNdES test (ratio of high-impact missense, nonsense and splicing mutations to low-impact missense (effectively-synonymous) and synonymous mutations) in cancer genes. The effectively synonymous mutations are missense mutations that have AlphaMissense pathogenicity score < 0.39. **b)** histogram of log10 of the dEXdIN (top panel) and dNdES (bottom panel) effect sizes in cancer genes. **c)** The slope of the linear regression between the dEXdIN and dNdES ratios at different AlphaMissense pathogenic scores, showing that range 0.35-0.57 results in slope approximately equal to 1.
